# PU.1 drives specification of pluripotent stem cell-derived endothelial cells to LSEC-like cells

**DOI:** 10.1101/2020.08.10.244517

**Authors:** Jonathan De Smedt, Elise Anne van Os, Irene Talon, Sreya Ghosh, Burak Toprakhisar, Rodrigo Furtado Madeiro Da Costa, Samantha Zaunz, Marta Aguirre Vazquez, Ruben Boon, Pieter Baatsen, Ayla Smout, Stefaan Verhulst, Leo A. van Grunsven, Catherine M. Verfaillie

**Affiliations:** Department of Development and Regeneration, Stem Cell Institute, KU Leuven, Leuven, Belgium; Liver Cell Biology research group, Vrije Universiteit Brussel (VUB), Brussels, Belgium; The Massachusetts General Hospital Cancer Center, Harvard Medical School, Boston, Massachusetts 02114, USA; The Broad Institute of Harvard and MIT, Cambridge, Massachusetts 02142, USA; Electron Microscopy Platform of VIB Bio Imaging Core at KU Leuven and VIB-KU Leuven Center for Brain & Disease Research, Leuven, Belgium

## Abstract

To date there is no representative *in vitro* model for liver sinusoidal endothelial cells (LSECs), as primary LSECs dedifferentiate very fast in culture and no combination of cytokines or growth factors can induce an LSEC fate in (pluripotent stem cell-derived) endothelial cells (ECs). Furthermore, the transcriptional programs driving an LSEC fate have not yet been described. Here, we first present a computational workflow (*CenTFinder*) that can identify transcription factors (TFs) that are crucial for modulating pathways involved in cell lineage specification. Using *CenTFinder*, we identified several novel LSEC-specific protein markers such as FCN2 and FCN3, which were validated by analysis of previously published single-cell RNAseq data. We also identified PU.1 (encoded by the *SPI1* gene) as a major regulator of LSEC-specific immune functions. We show that *SPI1* overexpression (combined with the general EC transcription factor *ETV2*) in human pluripotent stem cells (PSCs) induces ECs with an LSEC-like phenotype. The ETV2-SPI1-ECs display increased expression of LSEC markers such as CD32B and MRC1 as well as several of the proposed novel markers. More importantly, ETV2-SPI1-ECs acquire LSEC functions, including uptake of FSA-FITC as well as labelled IgG. In conclusion, we present the *CenTFinder* computational tool to identify key regulatory TFs within specific pathways, in this work pathways of lineage specification, and we demonstrate its use by the identification and validation of PU.1 as a master regulator for LSEC fating.

## Introduction

Liver sinusoidal endothelial cells (LSECs) are highly specialised endothelial cells (ECs) lining the sinusoidal capillaries of the liver where they reside along the space of Disse separating them from hepatocytes. LSECs have five main functions. First, LSEC fenestrations actively regulate the flow of macromolecules such as lipids and chylomicron remnants towards the hepatocytes (1,2). Second, LSECs endocytose larger lipid complexes, soluble macromolecules, hyaluronan, and glycosylation end products via the mannose receptor (MR or CD206), CD32B, stabilin-1 (STAB1), and stabilin-2 (STAB2) (3,4). Third, in response to hepatic injury, LSECs regulate liver regeneration by releasing hepatocyte growth factor (HGF), WNT2, and angiopoietin-2 (ANGPT2)(5,6). Fourth, LSECs can clear a multitude of viruses (7–12). Furthermore, LSECs clear bacterial compounds via the MR, STAB1, and STAB2 receptors. LSECs also express Toll-like receptors (TLRs) as well as several proteins involved in inflammasome formation (such as NLRP-1, NLRP-3, and AIM2)(13). Expression of CD32B allows uptake of IgG immunocomplexes while STAB2 is involved in endocytosis of gram-positive and -negative bacteria. Finally, LSECs have a major role in adaptive immunity. LSECs are antigen-presenting cells through MR-mediated antigen uptake and expression of MHC-I (14). Upon antigen presentation by MHC-I, LSECs can activate CD8+ T cells. Additionally, LSECs are the main cells in the liver responsible for the conversion of CD4+ T cells into CD4+ CD25+ Foxp3+ regulatory T cells (15).

In pathological conditions, LSECs become dysfunctional and play a role in non-alcoholic fatty liver disease (NAFLD) and fibrosis (16–19). In liver fibrosis, LSECs dedifferentiate, lose their fenestrations and deposit a basement membrane (a process called capillarisation). As a result, when chronic fibrosis or cirrhosis ensues, lack of blood flow towards hepatocytes aggravates hepatotoxicity and inflammation, leading to portal hypertension (20).

It should therefore be clear that LSECs, that play important roles in crosstalk with other (non-) parenchymal liver cell types in health and disease, should be included in liver co-culture models. However, primary LSECs very quickly lose their characteristics (e.g. fenestrae and endocytic capacity) and undergo apoptosis within 4-6 days of *in vitro* culture (21–23). Furthermore, limited availability of human primary LSECs is a major obstacle in the progression of LSEC research. A few immortalised LSEC lines exist, such as SK-HEP-1, that retain some LSEC characteristics (24). However, SK-HEP-1 cells do not take up FITC-ALB (24) and immortalised lines may not respond similarly in co-culture and disease models as primary LSECs do.

A number of protocols have been described to differentiate pluripotent stem cells (PSCs) towards ECs (25–27). However, only a few studies have attempted to differentiate PSCs into LSECs, either through culturing pluripotent stem cell-derived ECs in hypoxic conditions with a TGFβ inhibitor (28), or the combination of cAMP and a TGFβ inhibitor (29). Classically, differentiation protocols for a given cell type deploy sequential growth factor and cytokine cocktails to mimic *in vivo* differentiation. However, relatively little is known regarding LSEC development and, hence, signals required for LSEC differentiation *in vitro*. It has been hypothesised that LSECs derive from the septum transversum mesenchyme (30–32). However, recent lineage tracing experiments suggest that a large portion of the liver vasculature may derive from the sinus venosus endocardium (33). It is yet to be elucidated, though, whether the endocardium contributes to LSECs rather than macrovascular ECs. As an alternative, transcription factors (TFs) have been overexpressed to guide differentiation of PSCs towards cell types of all germ layers (34–37). However, the transcriptional programs or TFs that drive LSEC development are not well known.

Here, we designed a novel integrative meta-analysis pipeline (available as the *CenTFinder* R package) to facilitate analysis of published EC microarray data and identified pathways and TFs that might be crucial in fating LSECs. We combined several complementary analysis methods, including gene co-expression analysis (using WGCNA (38)), TF motif binding enrichment analysis (using RcisTarget (39)), and differential expression analysis on bulk RNAseq data (using DeSeq2 (40)), and identified several candidate TFs that may play a role in LSEC differentiation. As *SPI1*, encoding for PU.1, was identified as one of the top candidate TFs central to transcriptional networks for LSECs, we overexpressed *SPI1* in PSCs, and demonstrated that this drives specification of PSC progeny to an LSEC-like phenotype.

## Materials and methods

### Microarray selection and retrieval

We defined query key words, i.e. “endothelial cell AND Homo sapiens” and “LSEC AND Homo sapiens”, to construct URLs with base *https://www.ebi.ac.uk/arrayexpress/xml/v3/experiments?keywords=* to query the ArrayExpress database using the *RCurl* R package. ArrayExpress study accession codes were extracted from the HTTP responses. Subsequently, the respective SDRF files were downloaded to select relevant samples within each study. Only Affymetrix samples that were of human non-cancerous origin, and that were biotin-labelled were kept for analysis. Subsequently, new URLs were constructed to download the selected CEL files.

### Microarray raw analysis

Platform information was extracted from all downloaded CEL files using the *affyio* R package. Microarray platforms with only one downloaded CEL file were excluded from analysis. Subsequently, for each platform type, the respective CEL files were RMA normalised using the *oligo* R package. Probe intensity values were aggregated per gene by their geometric mean.

### Platform selection and platform effect removal

As microarray platforms differed in the number of genes for which probes were coated, we weighed the benefits of analysing more samples and hence fewer genes in common or analysing fewer samples with more genes in common. For standard analyses, we selected as a rule of thumb those platforms for which the product of the number of samples and the number of common genes was maximised. However, the LSEC arrays of platform type HG-U219 contained probes for a slightly smaller than optimal number of genes. Nevertheless, as these were the only retrieved LSEC arrays, we included these arrays as well and, hence, the downstream analysis was performed with 279 EC arrays and 16,248 genes. Subsequently, we applied the Combat algorithm of the *sva* R package to remove platform and batch confounding effects.

### Gene filtering

We excluded TFs that did not change by at least two-fold across all EC microarray samples, as such minimal expression changes are not likely to be biologically meaningful. A list of human TFs was downloaded from http://humantfs.ccbr.utoronto.ca. Furthermore, to save on computational time, we restricted the number of genes that were not TFs to the 7 551 most variable ones (log2 fold change standard deviation > 0.5961016). The total number of genes for downstream analysis was thus set to 9 000.

### Weighted Gene Correlation Network Analysis (WGCNA)

WGCNA was performed using the *WGCNA* R package. We constructed a signed network and tested soft-thresholding powers ranging from 1 to 30. A cluster dendrogram was created based on the topological overlap matrix with a minimum cluster size of 30. Subsequently, modules with a dissimilarity less than 0.3 were merged.

### Gene Ontology and KEGG pathway enrichment analysis

Gene Ontology enrichment was calculated for all identified WGCNA modules, using Fisher’s exact test from the *topGO* R package. The top 15 GO terms were retained for each GO category (i.e. Biological Process, Cellular Component, and Molecular Function).

### RcisTarget analysis

We imported motif rankings from the ‘*h19-tss-centered-10kb-7species*.*mc9nr*’ database (https://resources.aertslab.org/cistarget/). Overrepresented DNA binding motifs were identified within each of the WGCNA modules using the *RcisTarget* R package according to the creators’ guidelines.

### Bulk RNA sequencing analysis

#### Sample acquisition

PSCs, wherein the coding region of the TF *ETV2* (controlled by a TET-ON system) was recombined in the *AAVS1* locus (as described in Ordovas et al., 2015 (41)), were differentiated until day 10. RNA samples (N=3) were prepared and sequenced on an Illumina NextSeq 500 platform by the VIB Nucleomics Core (KULeuven, Belgium). Three RNAseq samples annotated as primary human LSECs were obtained from ArrayExpress (E-GEOD-40291 and E-GEOD-43984). However, only in one of these samples (E-GEOD-43984) LSEC specific genes, *FCGR2B, CLEC4M, STAB2*, and *CLEC4G*, were expressed. The other samples were excluded from analysis.

#### Read trimming

*Cutadapt* was used to trim the adapter sequence from all reads. Bases with Phred-scores lower than 20 were trimmed. Poly A {10} tails were removed and reads shorter than 20 bases were removed.

#### Genome index generation and read alignment

The human genome fasta files (GRCh38.92) and GTF file were downloaded from Ensemble, and assembled into a genome index using *STAR v020201*. Subsequently, *STAR* was used to align the filtered reads to the genome index. The R implementation of *FeatureCounts* (*Rsubread*) was used to generate raw read count matrices. Subsequently, we used the *GTFtools* Python package to calculate merged exon lengths, i.e. the union of all exons. Finally, we calculated the TPM matrix.

#### Differential gene expression analysis

The *DESeq2* R package was used to calculate differential gene expression (Benjamini-Hochberg corrected p-value<0.05). An absolute fold change of less than two was considered not biologically meaningful.

### Module Gene Set Variant Analysis

We used the modules identified by WGCNA as genesets in Gene Set Variant Analysis using the *GSVA* R package. Module gene sets were scored for each individual sample.Subsequently, mean GSVA scores were calculated per cell type to identify which modules were more prominently expressed for each cell type.

### Transcription factor ranking and visualisation

TFs were considered for overexpression if they were higher expressed in LSECs compared to endothelial cells derived from pluripotent stem cells through overexpression of *ETV2* (referred to as ETV2-ECs). Additionally, TFs were only retained if their kME centrality measure was higher than 0.5, and if their respective clusters were more active (GSVA) in LSECs than in other ECs. Next, the TFs were ranked based on the fraction of genes in their respective modules they regulated (inferred by RcisTarget). The modules of interest were visualised using Cytoscape. Regulons of the modules of interest were extracted and reformatted into a dataframe with Source and Target column, suitable for Cytoscape network import. TFs of the modules of interest were visualised when differentially expressed, regardless of whether they were higher or lower expressed in LSECs. For visualisation purposes we only retained the 20-most differentially expressed downstream targets, as evaluated by bulk RNAseq data. TFs assigned by WGCNA to other modules than the modules of interest were given the color of the module of interest if they had a high cluster centrality in the respective module of interest (kME>0.5). Node size was increased with increasing number of downstream targets.

### Single-cell RNA sequencing analysis

Fastq files from a recent human liver single-cell RNAsequencing study (GSE124395 (42)) were downloaded from the European Nucleotide Archive. Cellular barcode and UMI handling was done with UMI-tools. We used Cutadapt to trim bases with Phred-score < 20 and to remove reads shorter than 20 bases. Read quality was assessed with FastQC. STAR was used to create a human genome index based on the GRCh38.92 release of Ensemble. STAR was used as well for alignment. The count matrix was generated with UMI-tools. Cell quality control was performed using the *Scater* and *Seurat* R packages. Briefly, we removed cells with less than 2500 UMIs or less than 1500 detected genes. Only cells with a mitochondrial read content less than 15% were included for analysis. Only genes with a count higher than one in at least two cells were included for analysis. ENHANCE (43) was used for imputation and denoising. The *scran* implementation (cyclone) of the method described by Scialdone et al. (44) was used for cell cycle inference. SCENIC (with GRNBoost2 for ensemble learning) was used for detection of active gene regulatory networks (39). *Seurat* was used to correct for confounding variables, i.e. number of detected genes, number of total counts, cell cycle status, and percentage of mitochondrial counts.

The count matrix of a second study (GSE115469 (45)) was retrieved from the Gene Expression Omnibus. For each study, clusters were identified by Seurat and cell types were inferred by manually evaluating marker genes within the different clusters.

ECs were defined based on higher levels of *CDH5* and *KDR* expression and LSECs based on higher expression of *CLEC4G, STAB1*, and *PECAM1. FCGR2B-low* and *FCGR2B-high* LSEC subsets could be distinguished. Differential expression analysis was performed with a Wilcoxon rank sum test followed by a Benjamini-Hochberg multiple-test correction.

### Stem cell culture

H9 embryonic stem cells (ESCs) (WA09) were cultured in E8 Flex medium (cat. A2858501, ThermoFisher Scientific). Cells were passaged at 75% confluency using 0.1% EDTA in PBS (and tested for mycoplasma).

### Genome engineering of ESC lines for endothelial differentiation

Two million H9 ESCs, engineered with an FRT and FRT3 site in the *AAVS1* safe harbour locus (41), were nucleoporated with a plasmid mix containing 2µg of pCAGGS-FLPe (MES4488, Open Biosystems;without the puromycin resistance gene) and 8µg of the donor plasmid containing the gene of interest. Donor plasmids also contained FRT and FRT3 sites in identical orientation, a promotorless puromycin cassette for gene trapping, and an inducible TETon system for overexpression of the gene of interest. The coding sequences of NM_014209.3 and of NM_001080547.1 were cloned into the donor plasmid template for overexpression of *ETV2* and *SPI1* respectively. *ETV2* and *SPI1* coding sequences were linked by a P2A autocleavage sequence. Nucleoporation and subsequent selection was performed as described by Ordovas et al., 2015 (41).

### Differentiation of ESCs into endothelial cells

H9 ESCs were passaged at a 1:6 ratio and grown for 1-2 days in E8 Flex medium to about 40% confluency. On day 0 of endothelial differentiation, medium was changed to liver differentiation medium (LDM) (composition as described in (37)) and 5µg/ml doxycycline. On day 2 of differentiation cells were grown in LDM with 5µg/ml doxycycline and 2% FBS. Cells were passaged on day 4, day 8, and day 12 in a 1:3 ratio. After day 12 no further passaging was performed. When 24-well plate formats were required, ECs were passaged on day 4 from 6-well plates into 24-well plates (1 well into 10 wells, equivalent to a 1:2 split) and no further passaging was done until the time of read-out on day 12 of differentiation. To passage ECs, cells were washed with PBS and detached with StemPro™ Accutase™ Cell Dissociation Reagent (Gibco™, A1110501) for 35 seconds at 37°C. Subsequently, accutase was removed, fresh medium added, and cells gently detached with a cell scraper. Endothelial cells derived from H9-ETV2 PSCs and H9-ETV2-SPI1 PSCs are further referred to as ETV2-ECs and ETV2-SPI1-ECs respectively. In the conditions where cells were supplemented with VEGFA (50ng/ml) (cat. 100-20, Peprotech), it was added to the culture medium as of day 6 of differentiation **(Supplementary Figure 1)**. (N=7 for samples without VEGFA, N=4 for samples with VEGFA; statistical differences assessed by two-sided Student’s T-test)

**Figure 1.**
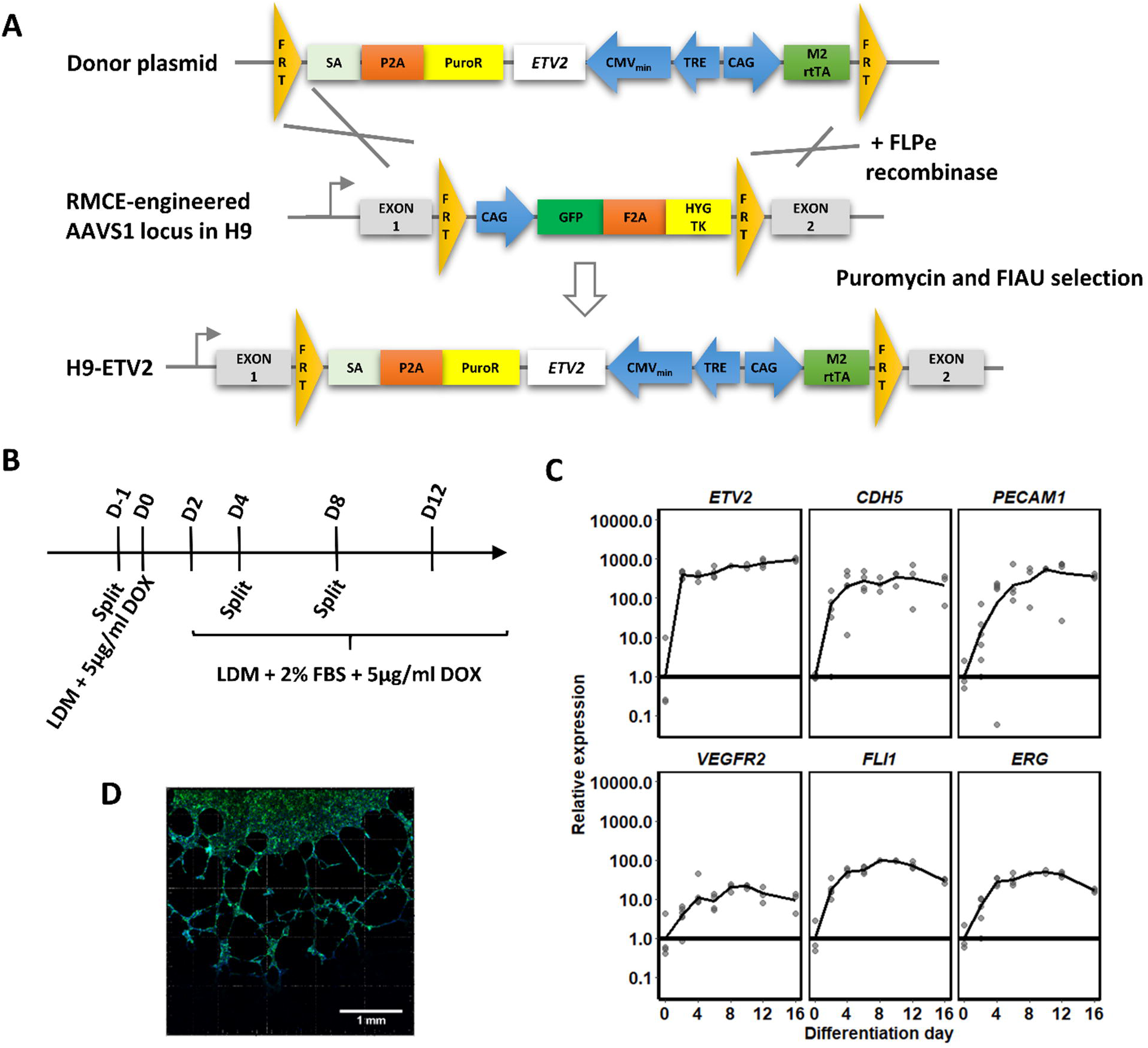
Generation and characterisation of ETV2-ECs. **A**. H9 human embryonic stem cells engineered with an RMCE cassette were recombined with a doxycycline inducible *ETV2* overexpression cassette (method described in and figure adapted from Ordovas et al. (41)). (FRT: Flippase recognition target; SA: splice acceptor; PuroR: Puromycin resistance gene; TRE: tetracycline response element; M2rtTA: M2 reverse tetracycline transactivator.) **B**. Schematic representation for PSC differentiation into endothelial cells. **C**. Gene expression (RT-qPCR) profile of *ETV2*, the endothelial markers *CDHS, PECAM1*, and *VEGFR2*, and the endothelial TFs *FLI1* and *ERG*. Samples were collected every 2 to 4 days of differentiation until day 30 with N=3 biological replicates per time point. Expression is relative to that of day 0 of differentiation (i.e. stem cells). **D**. Confocal image of phalloidin-stained ETV2-ECs in Matrigel showing tube formation.

### Isolation of mouse LSECs and hepatic stellate cells

Primary liver cells were isolated from 10-to 25-week-old BALB/c mice, (N=3), as previously described (46). LSECs and hepatic stellate cells (HSCs) were plated on collagen-coated wells in a 24 well plate (200,000 cells/well) in 0% FBS-DMEM for LSECs and 10% FBS-DMEM for HSCs. All methods, experimental protocols and animal experimentation ethics were carried out in accordance with the approved guidelines of the Vrije Universiteit Brussel (VUB, Belgium) and according to European Guidelines for the Care and Use of Laboratory Animals. All animal experimentation protocols were approved by the Ethical Committee of Animal Experimentation of the Vrije Universiteit Brussel (VUB, Belgium) (LA 123 02 12, project 18-212-1).

### Lentiviral overexpression of *SPI1, ERG*, and *FLI1*

H9-ETV2 PSCs were differentiated into ECs and passaged into 24-well plates on day 4. On day 6, cells were transduced with lentiviral vectors encoding for *SPI1, FLI1*, and *ERG1* (for *ERG* overexpressions: N=2; all other conditions: N=3) in the presene of 4µg/ml polybrene (cat. TR-1003-G, Merck) **(Supplementary Figure 2)**. Where *ETV2* overexpression was substituted for lentiviral overexpression of *FLI1* and/or *ERG*, doxycycline was omitted from culture media from day 8 of differentiation. Statistical differences were assessed by a two-sided Student’s T-test.

**Figure 2.**
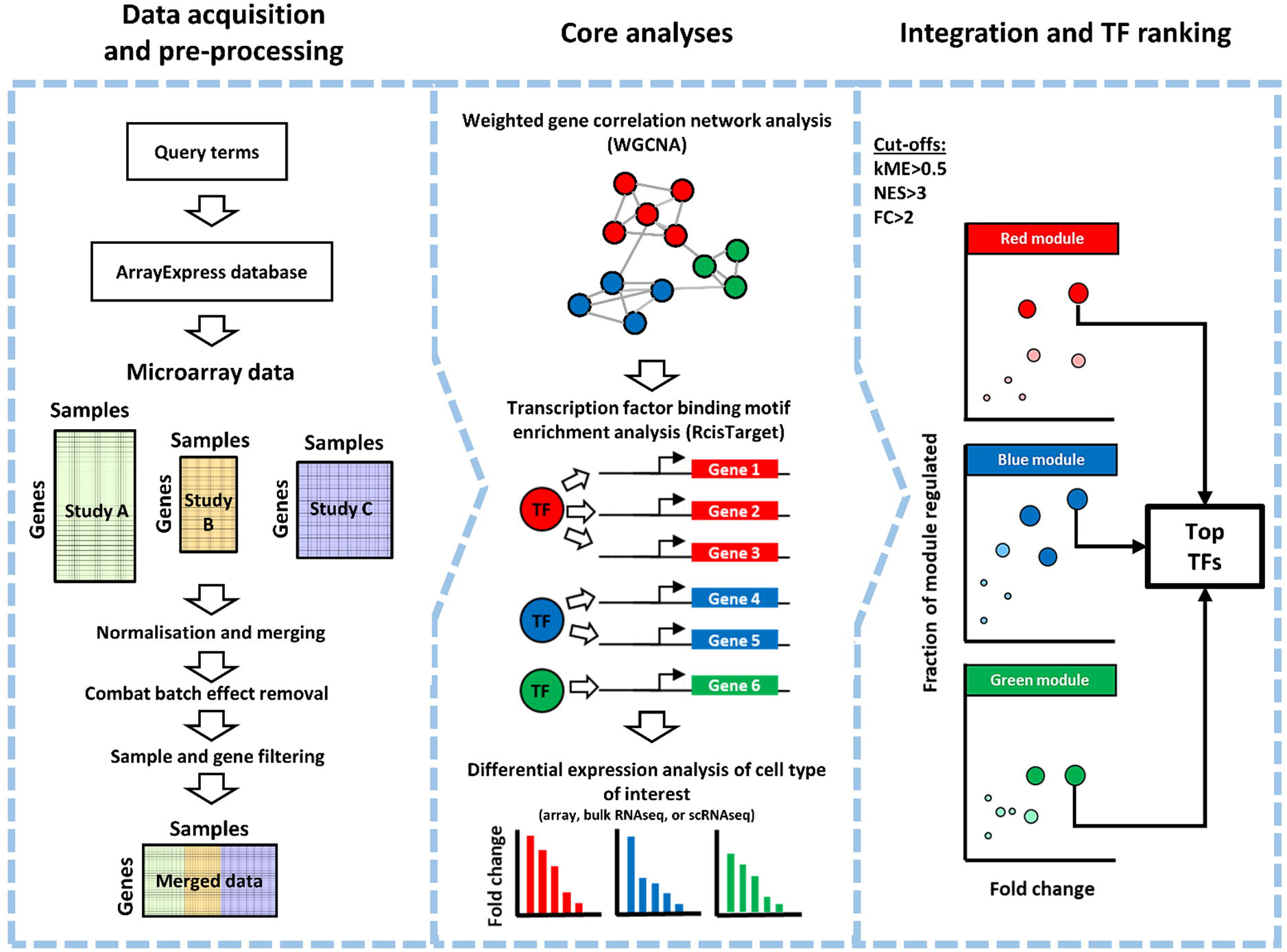
CenTFinder computational workflow. Schematic overview of the CenTFinder workflow for identification of TFs (central in differentiation and specification). **Left panel: Data acquisition and pre-processing**. The analysis starts by defining queries that are used for filtering and retrieving relevant expression data from the ArrayExpress database. Retrieved microarray data (of different formats) are combined and normalised and batch corrections are applied. Alternatively, a pre-processed expression matrix can be provided as input. **Middle panel: Core analyses**. The expression matrix is subsequently subjected to gene co-expression analysis (WGCNA), TF binding motif enrichment analysis (RcisTarget), and the results are overlapped with differential expression analysis. **Right panel: Integration and TF ranking**. The last step in the workflow is to combine the output of WGCNA, RcisTarget, and differential expression analyses to prioritise and rank TFs. Additional downstream analyses, e.g. Gene Ontology enrichment, and Cytoscape network visualisations are performed at this step.

### Reverse-transcriptase quantitative polymerase chain reaction

Cells were lysed and RNA extracted with the GenElute™ Mammalian RNA Extraction Kit (cat. RTN70, Sigma-Aldrich) and reverse transcribed into cDNA using SuperScript® III First-Strand Synthesis SuperMix (cat. 11752050, Invitrogen™), according to the manufacturer’s protocol. RT-qPCR reactions were made using the Platinum® SYBR® Green RT-qPCR SuperMix-UDG kit (cat. 11733046, ThermoFisher Scientific). Reactions were denatured at 95°C for 20 sec, followed by 40 cycles of a 1 sec denaturation step at 95°C and a 20 sec annealing and elongation step at 60°C. Gene expression values were normalised for *GAPDH, RPL19, RPS23*, and *EEF1A1*. Primer sequences are listed in Supplementary Table 1.

### Immunostaining

Culture medium was removed and cells (on coverslips) washed with PBS. Cells were fixed with 4% paraformaldehyde for 10 minutes, washed with PBS and permeabilised with 0.2% Triton X-100 in PBS. For CD32B staining, cells were not permeabilised. Blocking was performed using 5% donkey serum in 0.2% Triton X-100 in PBS. Primary antibodies (Supplementary Table 2) were diluted according to manufacturer’s guidelines in DAKO Antibody Diluent (cat. S202230, Agilent). Cells were incubated with the primary antibody mixes overnight at 4°C, washed with 0.2% Triton X-100 in PBS, and incubated for 30 minutes at room temperature with the secondary antibody (Supplementary Table 2)(1:500 diluted in DAKO Antibody Diluent). Cells were washed with 0.2% Triton X-100 in PBS, mounted with Prolong Gold (cat. P-36931, ThermoFisher Scientific), and visualised using a Zeiss Axioimager microscope. Stainings were replicated three times.

### Flow cytometry

Day 12 PSC progeny were detached with accutase for 35 seconds at 37°C. Accutase was inactivated by incubating cells in LDM + 10% FBS for 6 minutes at room temperature. Cells were washed with PBS and resuspended in PBS with 1% BSA (cat. A7979-50ml, Sigma Aldrich). Cells were incubated with primary antibodies (Supplementary Table 2) for 30 minutes at 4°C, washed with PBS with 1% BSA and incubated with secondary antibodies (Supplementary Table 2) for 30 minutes at 4°C. Viability was assessed by propidium iodide (cat. 81845-25MG, Sigma Aldrich). Flow cytometry was performed using a BD FACSCanto™ II High Throughput Sampler. Cells were gated according to the strategy depicted in **Supplementary Figure 3**. Two-sided paired Student’s T-tests were applied on population median intensities (replicates: N = 15 for CD32B and N=14 for MRC1).

**Figure 3.**
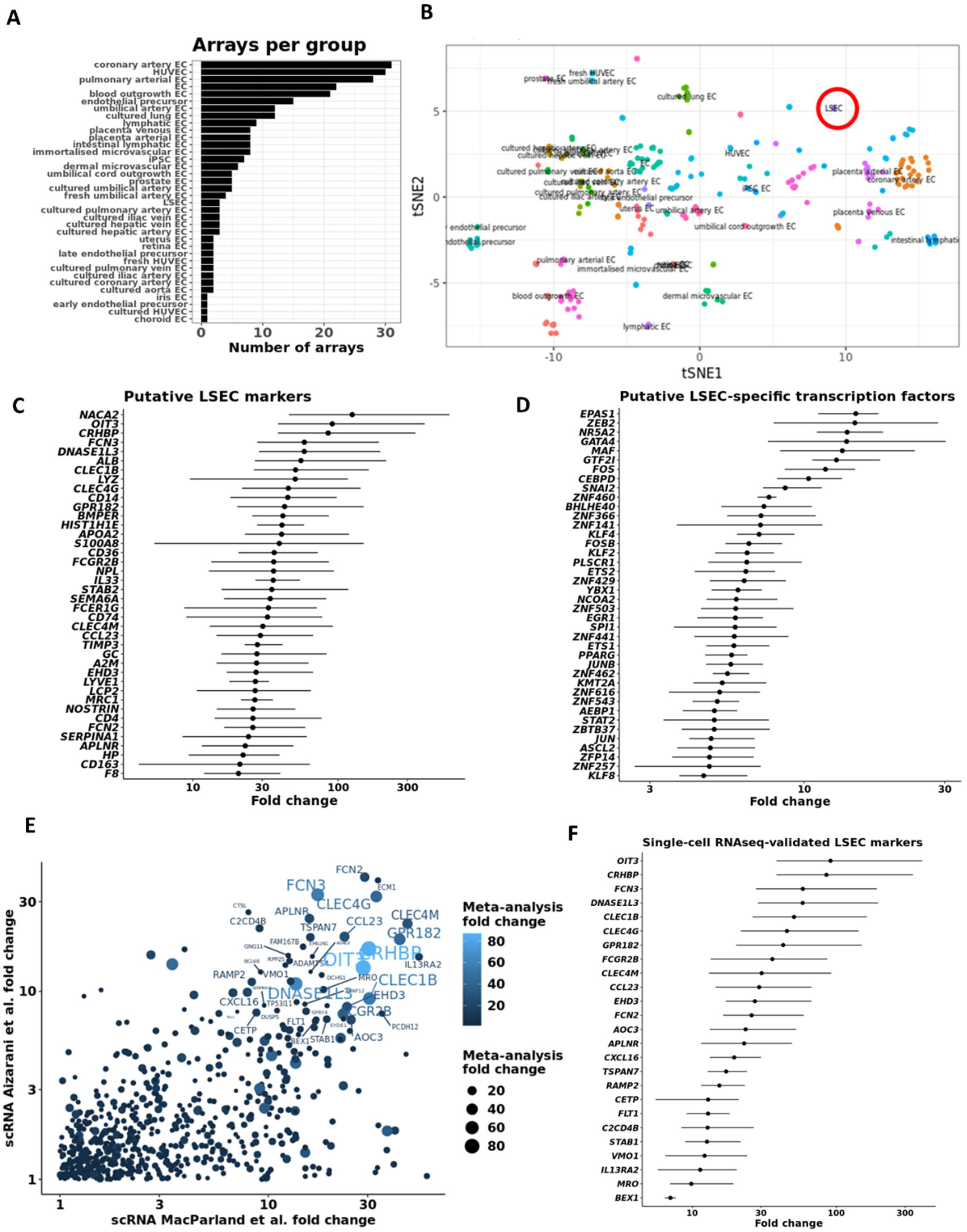
Intersection between microarray meta-analysis and single-cell RNAseq identifies putative LSEC markers. **A**. Bar chart indicating the number of microarrays per endothelial cell type that was included in the CenTFinder analysis (total number of arrays = 279). **B**. tSNE representation of all endothelial cells used in the analysis (LSECs circled in red). LSEC microarray data were contrasted to the expression data of all other endothelial cells used in the meta-analysis. Line charts show putative markers **(C)** and TFs **(D)** with increased expression in LSECs compared to all other endothelial cells (assessed by Wilcoxon rank sum test). Putative markers and TFs were sorted by fold change. Dots indicate the mean fold change between LSECs and all other ECs. Lines span the interquartile range. **E**. Overlap of genes that are specifically expressed in LSECs in the single-cell RNA sequencing studies and in the microarray meta-analysis. The identified genes are more expressed in LSECs compared to all other endothelial cells as well as compared to all other parenchymal and non-parenchymal liver cells. X- and Y-axes denote the two single-cell RNAseq studies (42, 45). **F**. LSEC marker ranking by fold change compared to all other ECs in the microarray meta-analysis. For this analysis, genes were ranked only if in each of the studies they were more than 7-fold higher expressed in LSECs compared to the other cells. Dots indicate the mean fold change between LSECs and all other ECs. Lines span the interquartile range.

### Tube formation assay

Wells of 24-well plates were coated with undiluted Matrigel® hESC-Qualified Matrix (protein concentration approximates 10mg/ml)(cat. 354277, Corning®). Matrigel was solidified by incubation for at least 10 minutes at 37°C. Day 12-ECs (H9-ETV2 and H9-ETV2-SPI1) were seeded on the matrigel at a density of 150,000 cells per well. After 20 hours, cells were fixed with 4% paraformaldehyde, washed with PBS and permeabilised with 0.2% Triton X-100 in PBS for 30 mins. Cells were washed with PBS and blocked with 0.1M glycine in PBS for 30 minutes. Subsequently, cells were incubated with 2 U/ml phalloidin (Supplementary Table 2), 2.5µg/ml DAPI (Supplementary Table 2) and 1% BSA in PBS for 90 minutes in the dark at room temperature. Cells were washed with PBS and tubes visualised by confocal imaging with a Zeiss LSM 880 – Airyscan.

### Scanning electron microscopy

ETV2-ECs and ETV2-SPI1-ECs were cultured on coverslips until day 12 of differentiation with VEGFA (50ng/ml) (cat. 100-20, Peprotech). BALB/c LSECs were cultured for 1.5-2 hours after isolation, with (40ng/ml, cat.V4512-10UG, Sigma Aldrich). Cells were washed thoroughly with PBS to avoid protein and serum remnants. Next, cells were immersed in 500µl PBS and fixed for 10 minutes with 500µl of 5% glutaraldehyde in 66mM cacodylate buffer. The fixative solution was removed and cells maintained overnight at 4°C in 2.5% glutaraldehyde in 66mM cacodylate buffer. Cells were washed with 0.1M cacodylate buffer at room temperature and post-fixed for two hours at room temperature with 1% OsO_4_ + 1.5% K_4_Fe(CN)_6_ in 0.1M cacodylate buffer. After post-fixation cells were washed with 0.1M cacodylate buffer and dehydrated with increasing ethanol concentrations (30% - 50% - 70% - 90% - 100% - 100%) with 5 minutes incubation per concentration. Finally, the samples were critical point dried in a Leica CPD300 critical point dryer and the coverslips with cells mounted on pin stubs with carbon stickers. Cells were observed and imaged in a Zeiss Sigma scanning electron microscope at an accelerating voltage of 5 kV. Pore diameters were counted and measured using ImageJ and the EBImage R package.

### FSA-FITC uptake assay

ETV2-ECs and ETV2-SPI1-ECs were cultured until day 12 of differentiation on Matrigel-coated wells in a 24-well plate. BALB/c LSECs were isolated and plated as well on collagen at 200.000 cells per well in a 24-well plate in 0% FBS-DMEM. After one hour of LSEC plating, ETV2-ECs, ETV2-SPI1-ECs, and LSECs were supplemented with 10µg/ml FSA-FITC (prepared as described in (47)) and 1:1000 IncuCyte® NucLight Rapid Red Reagent (cat. 4717, Sartorius). Cells were imaged hourly over a period of 24 hours using the IncuCyte Cell analysis system (Essen Bioscience). Quantification was performed with the IncuCyte Cell analysis software. This experiment was repeated twice. Statistical differences were assessed by mixed ANOVA.

### IgG-AF555 uptake assay

H9-ETV2 and H9-ETV2-SPI1 PSCs were differentiated to ECs as described above. On day 12 of differentiation cells were washed with PBS and incubated for 2 hours at 37°C with 100µg/ml IgG (conjugated with Alexa Fluor 555)(Supplementary Table 2) in LDM + 2% FBS + 5µg/ml doxycycline. Cells were washed with PBS and fixed with 4% paraformaldehyde. For imaging intended for statistical assessment of uptake, samples were additionally stained with DAPI. CellProfiler version 3.1.9 (48) was used for segmentation of the red channels using the Watershed algorithm. Mean intensities were calculated per object. For confocal imaging, samples were additionally stained with DAPI, phalloidin, and an anti-RAB5 antibody to detect endosomes (Details in Supplementary Table 2). This experiment was repeated twice. Co-localisation of RAB5 and IgG-AF555 was quantified using the JaCoP ImageJ plug-in, as described (49). The Pearson’s correlation coefficient was computed after thresholding.

### Statistical analysis of RT-qPCR

For all pair-wise comparisons (e.g. in the *FLI1* and *ERG* overexpression and VEGFA addition experiments) we applied a Student’s T-test. Statistical significance of the *SPI1* lentiviral titer gradient on *FCGR2B* and *LYVE1* expression was assessed using linear modelling. Time-course comparisons of RT-qPCR data and FITC-FSA were assessed by mixed ANOVA. Sample sizes were *a priori* decided through power analysis simulations in R, given a minimal effect size of interest of two cycle thresholds, an average standard deviation of one cycle threshold, a significance level of 0.05, and a minimally required power of 0.8.

### Code availability

All code is reproducible and can be accessed via https://github.com/jonathandesmedt92/CenTFinder.

## Results

### ETV2 overexpression fates pluripotent stem cells to the endothelial lineage

To effectively differentiate PSCs into ECs, we recombined a doxycycline-inducible *ETV2* overexpression cassette in H9 ESC already containing an FRT-flanked exchange cassette **(Figure 1A)** as described by Ordovas et al. (41)). Resulting ECs are further referred to as ETV2-ECs. We assessed the endothelial differentiation **(Figure 1B)** by RT-qPCR **(Figure 1C)**. *ETV2* was more than a 1000-fold overexpressed from day 2 of differentiation. As expected, expression levels of the pluripotency markers *NANOG* and *POU5F1* decreased by more than 90% within four days of differentiation **(Supplementary Figure 4A)**. On day 6 of differentiation expression levels of the endothelial markers, i.e. *PECAM1, CDH5*, and *VEGFR2*, and endothelial TFs, i.e. *FLI1* and *ERG*, were induced and remained stable **(Figure 1C)**. We confirmed expression of CD31 by immunostaining **(Supplementary Figure 4B)**. Low to no staining was observed for the LSEC markers CD32B and MRC1. Furthermore, tube formation, a key feature of ECs, was observed after seeding ETV2-ECs on matrigel **(Figure 1D)**.

**Figure 4.**
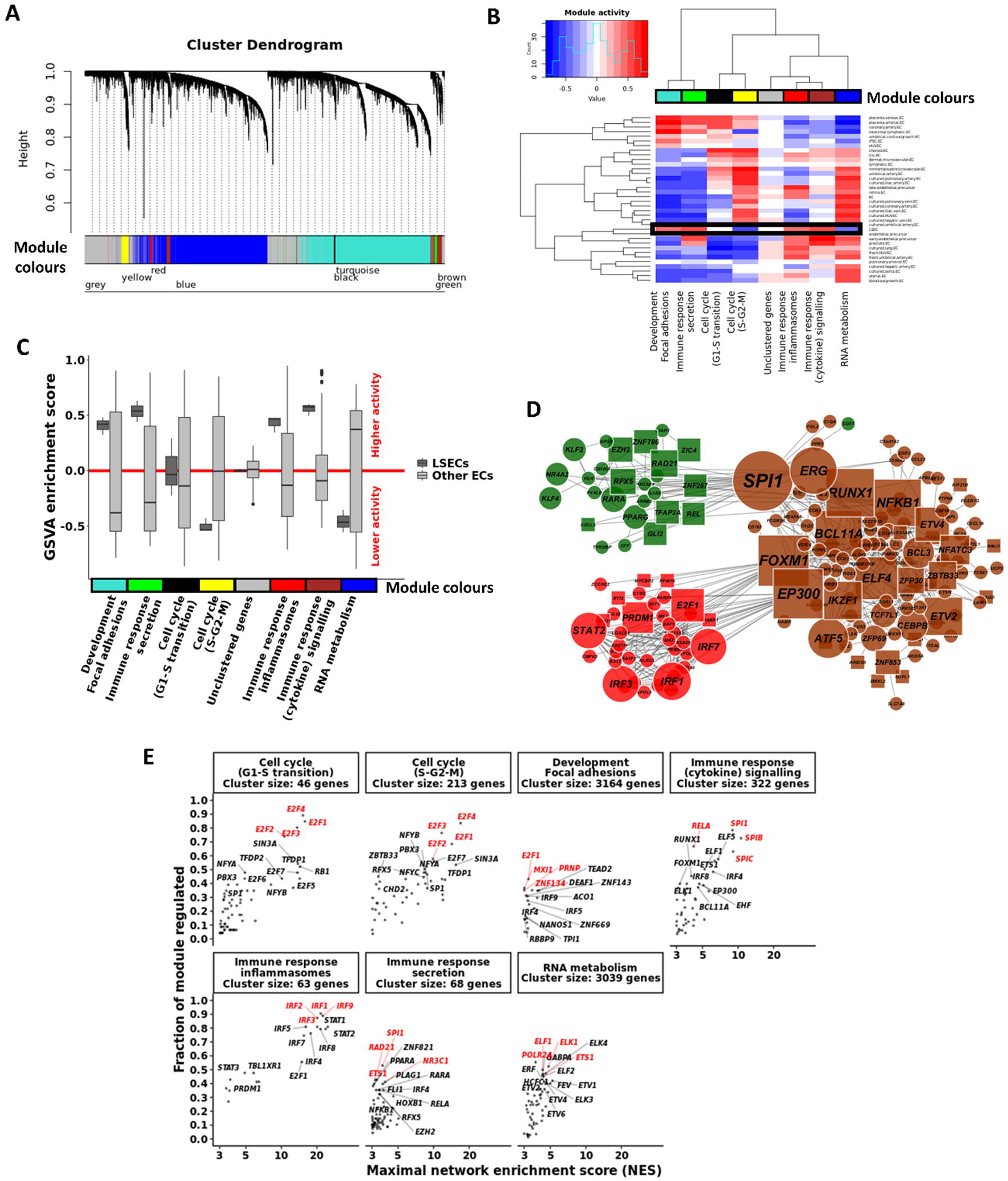
Gene co-expression (WGCNA) and transcription factor binding motif (RcisTarget) analyses. **A**. WGCNA cluster dendrogram visualising the gene co-expression modules identified in the microarray meta-analysis. These modules are active in at least one of the endothelial cell types. The grey module contains genes unassigned to any module due to low correlative values. **B**. Heatmap and clustering of the Gene Set Variant Analysis (GSVA) activity scores for each of the modules and for each of the cell types. **C**. Module colours correspond with those from the cluster dendrogram in A. Black box identifies module activities in LSECs. C. Boxplots of the GSVA enrichment scores for each of the modules, comparing LSECs to all other endothelial cell types. **D**. Cytoscape visualisation of the LSEC-specific immune response modules. Edges indicate a regulatory link identified by RcisTarget. TF node size relates to the number of downstream targets. For the sake of simplicity, only nodes of the 20 most differentially expressed downstream targets are shown for each TF. Nodes were coloured according to their module colour. TFs belonging to other modules but with a kME>0.5 for either the brown, red, or green modules were included as well and coloured accordingly. Circular (*square*) nodes represent genes that are higher (*lower*) expressed in LSECs compared to ETV2-ECs. **E**. Enrichment of TF binding motifs was calculated for each of the WGCNA gene co-expression modules. Each dot represents an enriched TF binding motif with the x-axis indicating the RcisTarget Network Enrichment Score (NES) and the y-axis indicating the fraction of genes in the respective module that has binding motifs for the respective gene. For genes with multiple enriched motifs, only the motif with the highest NES is shown. Top TFs are highlighted in red.

### CenTFinder workflow

To identify and rank TFs required for fating ETV2-ECs towards LSECs, we designed a computational strategy **(Figure 2)** to perform a meta-analysis on publicly available or in-house transcriptional data from various sources of ECs and LSECs (Supplementary Table 3). To ensure that the identified TFs play a ‘master regulator’ role in fating ECs to LSECs, we combined various computational methods such as gene co-expression, TF binding motif enrichment, as well as differential expression analyses. Aside from TF rankings for LSECs, we also constructed TF rankings (Supplementary Table 4) for all other EC types that were included in the meta-analysis (Supplementary Table 3). The entire workflow is available as the *CenTFinder* R package on Github (https://github.com/jonathandesmedt92/CenTFinder).

### Identification of LSEC markers by combination of microarray and single-cell RNAseq analyses

We downloaded, filtered, and normalised relevant microarray data **(Figure 2. Left pane** and **Figure 3A**, metadata in Supplementary Table 3). Next, we represented the gene expression profiles of all arrays (N=279) by t-distributed Stochastic Neighbor Embedding (t-SNE), which shows great heterogeneity and sets the human primary LSECs apart from all other EC subtypes **(Figure 3B)**. Differential expression analysis between LSECs and all other EC types **(Figure 3C-D)** confirmed established as well as only recently identified LSEC marker genes such as *CLEC4G, FCGR2B, STAB2, MRC1, CLEC4M, F8, LYVE1, FCN2, FCN3, OIT3, CLEC1B, DNASE1L3*, and *GPR182* (45,50,51). Additionally, we could identify putative novel LSEC marker candidates of which we validated LSEC-specific expression in two recently published liver scRNA seq data sets (42,45) **(Figure 3E)**. Several putative markers were only expressed in the LSEC population, i.e. *CDH5+ KDR+ CLEC4G+* cells (42,51). Additionally, some candidate markers, such as *ALB* and *APOA2*, were expressed in hepatocytes rather than LSECs. By merging fold changes of the *CenTFinder* and scRNAseq datasets, we generated a consensus list of scRNAseq-validated LSEC markers that were at least 7-fold higher expressed in LSECs compared to the other cells in the respective datasets (42,45) **(Figure 3F)**.

### WGCNA and RcisTarget analysis identified *SPI1* and *IRFs* as central to immune response transcriptional modules specific for LSECs

TFs were ranked by fold change (LSECs compared to other ECs in the microarray meta-analysis **(Figure 3D)**. However, such ranking identifies both putative TFs required for fating ECs to LSECs, as well as broadly acting TFs (e.g. *JUN* and *FOS*). Furthermore, transcripts of cell types other than LSECs may also bias this ranking as assessed by analysis of published single-cell RNAseq data (42) **(Supplementary Figure 5)**. We subsequently used a gene regulatory network (GRN) approach (Weighted Gene Co-expression Network Analysis; WGCNA) to select more specific TFs that might guide LSEC differentiation.

**Figure 5.**
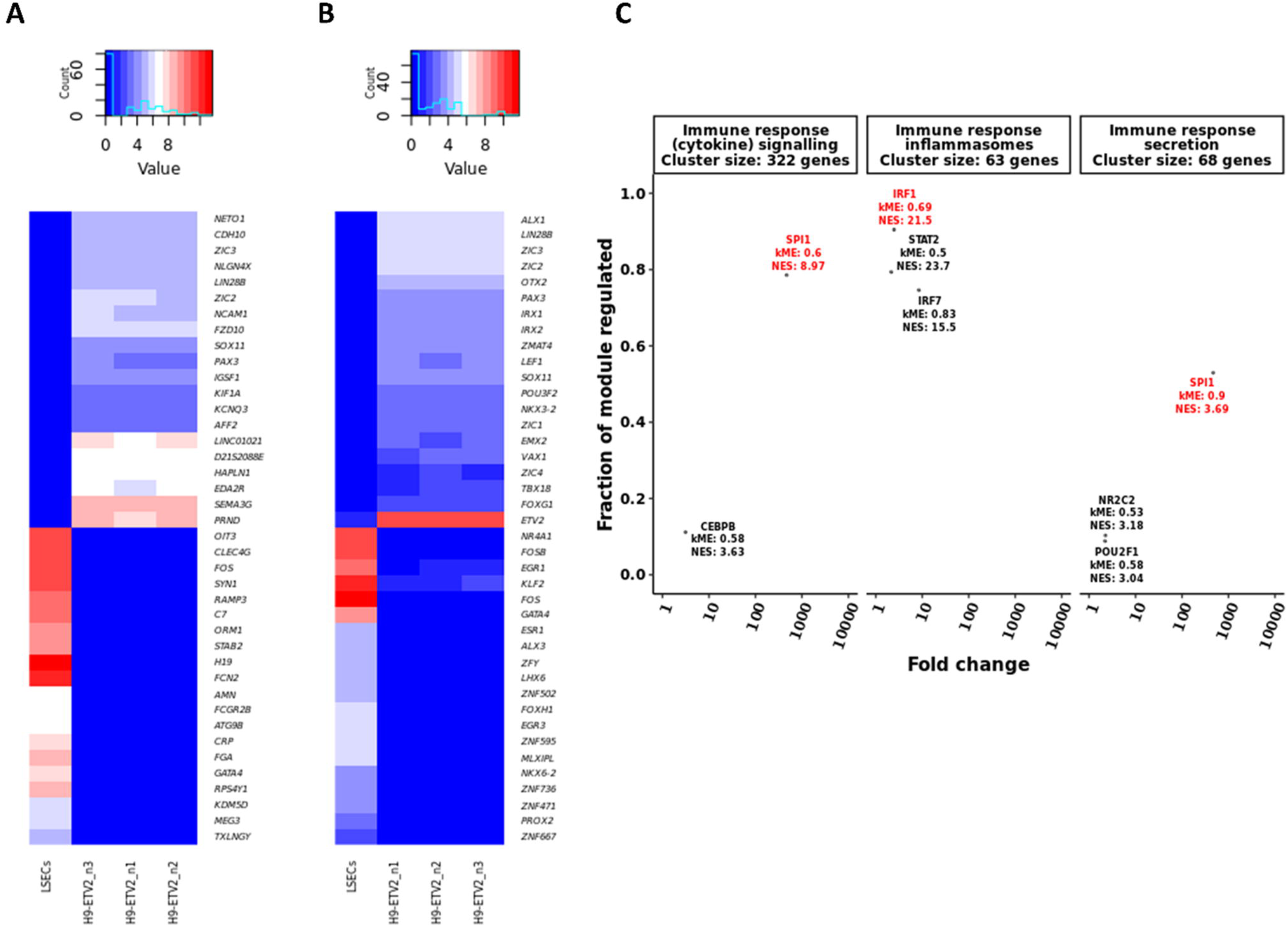
CenTFinder analysis identifies *SPI1* and *IRF* TFs as major LSEC regulators. **A**. Heatmap of the top 20 most up- and down regulated genes in LSECs compared to ETV2-ECs. **B**. Heatmap of the top 20 most up- and down regulated TFs in LSECs compared to ETV2-ECs. **C**. CenTFinder core analysis results were combined. Each dot represents a TF that was differentially and higher expressed in LSECs compared to ETV2-ECs. Additionally, only TFs with kME>0.5 and NES>3 were considered. *SPI1* was identified as the central TF in the immune response clusters related to cytokine signalling and secretion. *IRF* TFs were central to the immune response cluster related to inflammasomes. Top TFs are highlighted in red.

WGCNA is a tool that clusters genes with correlated expression patterns into modules that are, as a result, highly enriched for specific gene ontology (GO) terms and pathways. We subjected the expression matrix to WGCNA (38), to identify transcriptional modules, their associated GO terms and central TFs (i.e. TFs with a high module eigengene). We identified seven transcriptional modules **(Figure 4A)** that were each active in at least one of the EC types included in the meta-analysis **(Figure 4B)** (excluding the grey module containing unassigned genes). By GO analysis, three of the modules were related to immune response genes (i.e. red, green, and brown), two to cell cycle genes (i.e. (S-G2-M) yellow and (G1-S transition) black), one to RNA metabolism and translation (i.e. blue), and one to development, gene expression regulation, and focal adhesions (i.e. turquoise) **(Supplementary Figure 6)**. Comparison of module activities (calculated with Gene Set Variation Analysis, GSVA) (52) demonstrated that the modules related to immune response, development, and focal adhesions were more active in LSECs compared to other ECs. In contrast, the (S-G2-M) cell cycle and the RNA metabolism modules were less active in LSECS **(Figure 4B-C**. Interestingly, the immune clusters contained several C-type lectin receptors such as *CLEC10A, CLEC2D, CLEC4A, CLEC4E, CLEC5A*, and *CLEC7A*, as well as several immunoglobulin receptors such as *FCER1G, FCGR1A, FCGR2A, FCGR2B*, and *FCGR3B* (immune clusters represented in **Figure 4D**).

**Figure 6.**
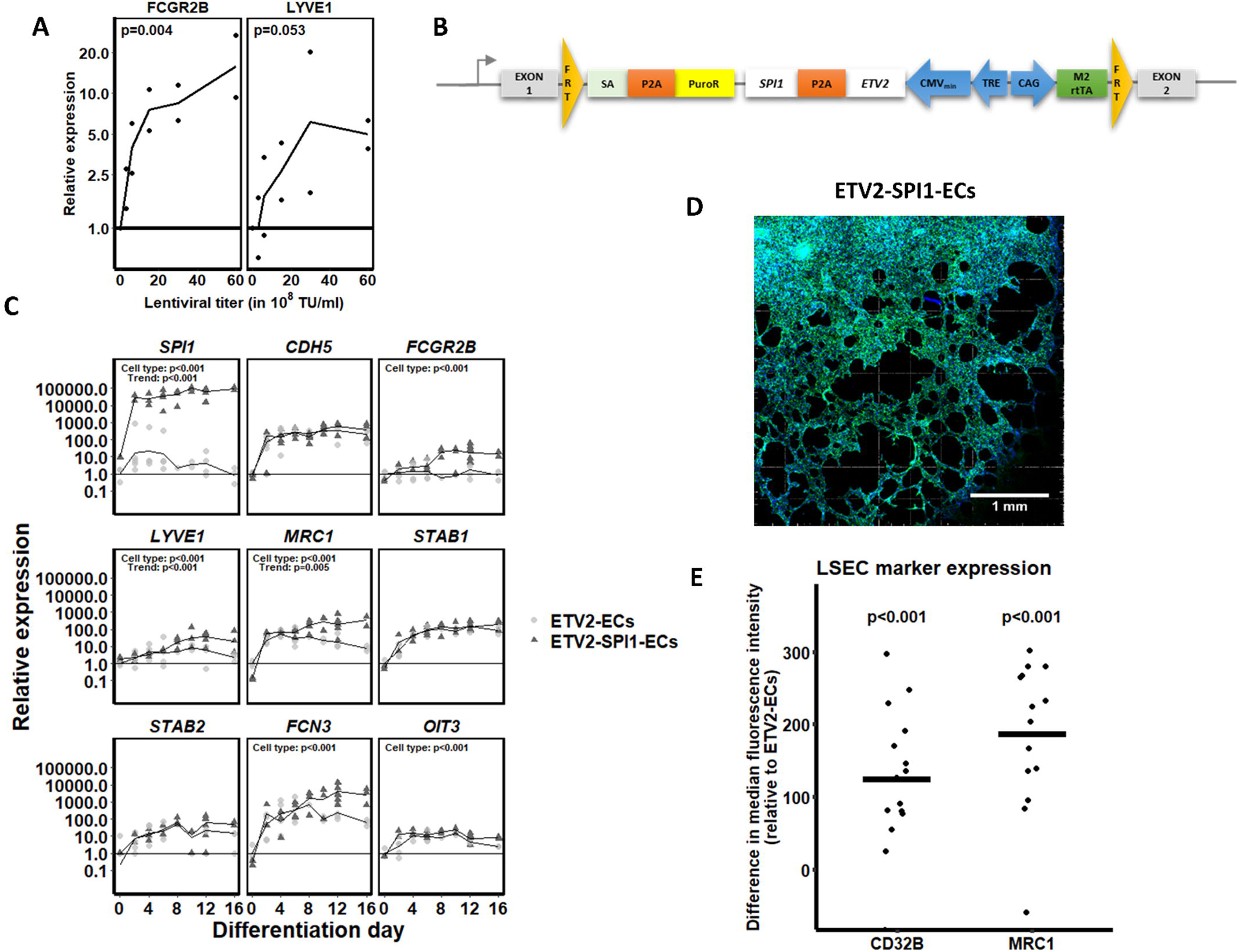
Characterisation of (liver sinusoidal) endothelial cell markers in ETV2-ECs and ETV2-SPl1-ECs. **A.** ETV2-ECs were transduced on day 6 of differentiation with increasing titres of the SP/1-encoding lentiviral vector. The effect of *SP/1* overexpression on the LSEC markers *FCGR28* and *LYVE1* is shown. Statistical analysis by linear modelling (Expression∼ Titre); N= 2 biological replicates per concentration. **B**. ETV2-SPl1 construct engineered into the *AAVS1* safe harbour locus. Recombination was performed analogously to Figure lA. (FRT: Flippase recognition target; SA: splice acceptor; PuroR: Puromycin resistance gene; TRE: tetracycline response element; M2rtTA: M2 reverse tetracycline transactivator). C. Time-course comparison between ETV2-ECs and ETV2-SPl1-ECs for the overexpressed *SPI1*, for the endothelial cell-specific gene *CDHS*, for the known LSEC markers *FCGR28, LYVE1, MRC1, STAB1*, and *STAB2*, and for novel LSEC markers such as *FCN3*, and *O/T3*. Statistical significances were assessed by mixed ANOVA. Undetected mRNA values were represented with a relative expression of 1. The x-axes denote the days of differentiation. The differentiation protocols for ETV2-ECs and ETV2-SPl1-ECs are identical. **D**. Tube formation assay on day 12 of differentiation of ETV2-SPl1-ECs. **E**. LSEC marker expression (CD32B and MRCl) in ETV2-SPl1-ECs compared to ETV2-ECs. Chart indicates the median fluorescence intensity shift for CD32B and MRCl of ETV2-ECs and ETV2-SPl1-ECs on day 12 of differentiation (replicates: **N** = 15 for CD32B and N=14 for MRCl). The CD32 and MRCl positive cells were gated from single-cell, Pl-negative, and VE-cadherin-positive populations (gating strategy in Supplementary Figure 3).

To avoid interference of spurious correlations due to technical or biological noise, we applied RcisTarget (39) to each of the WGCNA modules **(Figure 4E)**. RcisTarget identifies TFs for which there is an enriched binding motif presence in the cis-regulatory regions of the genes in a given gene set. Only TFs with a Network Enrichment Score (NES) higher than 3 were considered and TFs that regulate higher fractions of their respective modules were deemed more important or influential. In the immune response modules related to cytokine signalling and secretion, *SPI1* had binding motifs in most of the genes in each of the respective clusters. In the immune response module related to inflammasomes, most genes were regulated by *IRF* TFs.

To further narrow down more promising TFs, we used bulk RNA sequencing data to identify which of these TFs are differentially expressed between ETV2-ECs and LSECs **(Figure 5A-B)**. Next, we made an overlap between the cluster-central and RcisTarget-enriched TFs that were expressed more in LSECs compared to ETV2-ECs **(Figure 5C)**. All TFs in the development and focal adhesions module were less expressed in LSECs compared to ETV2-ECs. In the immune reponse (cytokine signalling and secretion) modules *SPI1* was the most highly differentially expressed (469-fold, p<0.001;cluster centrality kME=0.6 and kME=0.9 respectively, and network enrichment score NES=8.97 and NES=3.69 respectively). In the immune response (inflammasomes) module, *IRF1, IRF7*, and *STAT2* passed the filtering criteria.

In conclusion, the microarray EC and LSEC meta-analysis identified *SPI1* and *IRFs* to be candidate master regulators of LSEC fating. As the comparison of LSECs and ETV2-ECs also identified *SPI1* to be the highest ranking TF in LSECs, subsequent studies were done to validate *SPI1* as a master regulator of LSEC fating.

### *SPI1* overexpression induces LSEC marker expression

To evaluate if *SPI1* can drive LSEC specification and maturation in ETV2-ECs, we cloned the *SPI1* coding sequence into a pLVX backbone and transduced ETV2-ECs on day 6 of differentiation with the resulting lentiviral vector. Clear upregulation of the LSEC markers *FCGR2B* and *LYVE1* was observed **(Figure 6A)**. To avoid the toxicity associated with lentiviral transduction, and to ensure that all cells contained the *SPI1* TF, we next recombined a doxycycline-inducible cassette for both *ETV2* and *SPI1* in the *AAVS1* safe harbour locus, to generate ETV2-SPI1-ECs **(Figure 6B)**. Combined induction of *ETV2* and *SPI1* did not impact general EC differentiation **(Supplementary Figure 7)**. Specifically, expression of *NANOG* and *POU5F1* decreased by >90% on day 4 (**Supplementary Figure 7A**) and no differences in expression of the EC markers *PECAM1* and *CDH5*, and the EC TFs *ERG, ETV2*, and *FLI1*, were observed between ETV2- and ETV2-SPI1-ECs **(Supplementary Figure 7B)**. Consistently, immunostaining for CD31 was similar in ETV2 and ETV2-SPI-ECs **(Supplementary Figure 8)**. Transcript levels for *SPI1* were >1000 fold higher expressed in ETV2-SPI1-ECs compared to ETV2-ECs (p<0.001) **(Figure 6C)**. In addition, tube formation by ETV2-ECs and ETV2-SPI1-ECs was similar **(Figure 6D)**.

**Figure 7.**
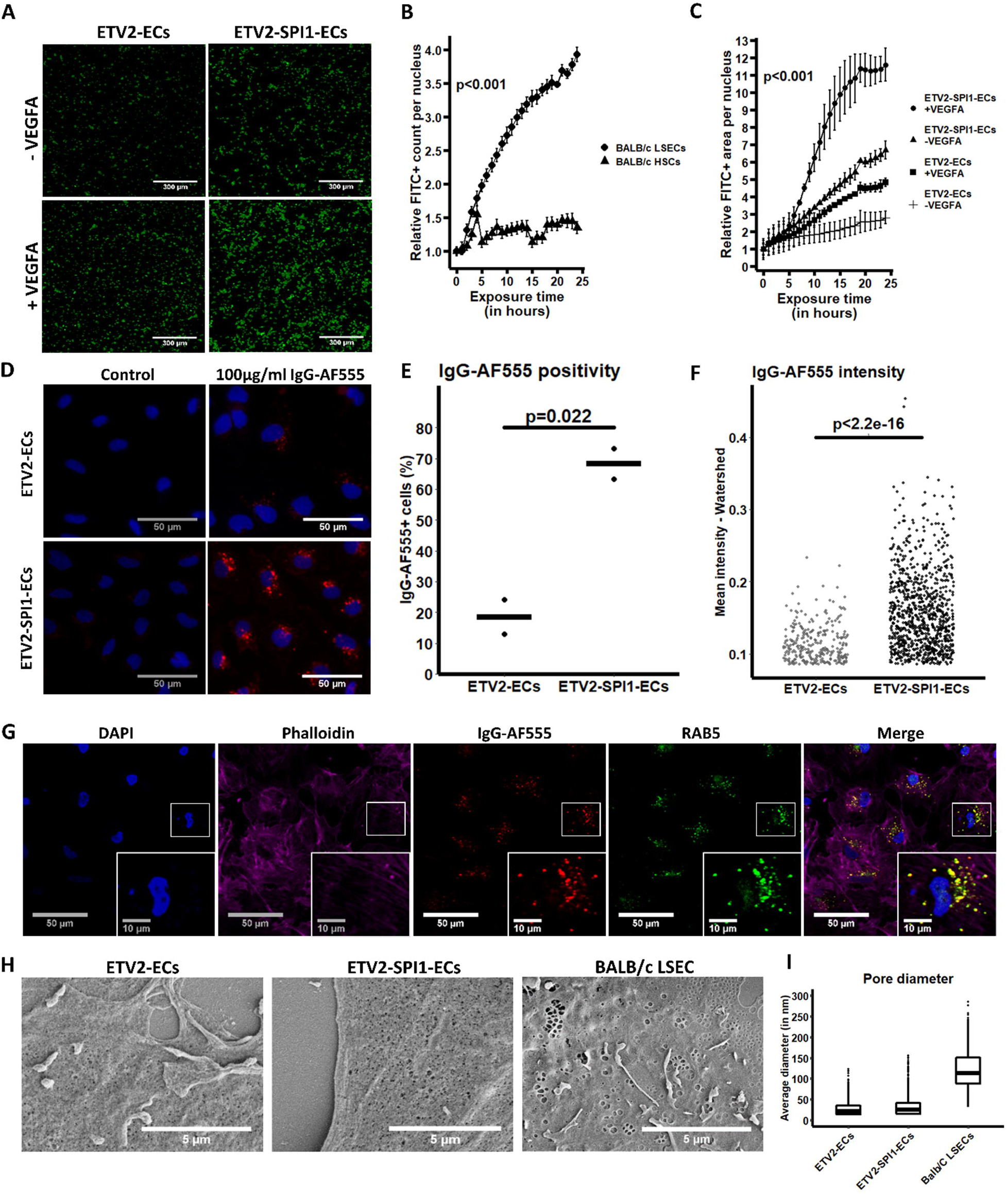
LSEC-specific functional characterisations of ETV2-ECs and ETV2-SPl1-ECs. **A-C**. FSA-FITC uptake assay (N=2 biological replicates) on ETV2-ECs and ETV2-SPl1-ECs with and without addition of VEGFA, as well as on BALB/c LSECs and hepatic stellate cells (HSCs). Images are representative for 24 hour exposure to FSA-FITC. Scale bars are 300µm. FITC fluorescence was measured every hour. Statistical differences were assessed by mixed ANOVA. **D**. lgG-AFSSS uptake assay. ETV2-ECs and ETV2-SPl1-ECs were exposed for two hours to l00µg/ml lgG-AFSSS to measure the clathrin-mediated endocytosis of lgGs upon their binding to CD32B. Scale bars are S0µm. (representative example of N = 3 biological replicates). **E**. Comparison of the percentage of lgG-AFSSS-positive cells in ETV2-ECs and ETV2-SPl1-ECs at day 12 of differentiation. Statistical assessment by Student’s T-test. **F**. Comparison of the AFSSS intensity of lgG-AFSSS-positive cells in ETV2-ECs and ETV2-SPl1-ECs at day 12 of differentiation. Statistical assessment by Wilcoxon rank sum test. **G**. Confocal imaging of ETV2-SPl1-ECs exposed to lgG-AFSSS. Cells were co-stained with an anti-RABS antibody to evaluate co-localisation of lgG-AFSSS and endosomes. **H**. Scanning electron microscopy of ETV2-ECs, ETV2-SPl1-ECs, and BALB/c LSECs. Scale bars are Sµm (representative example of all cells per coverslip). Average pore diameters are shown in **I**.

Next, we compared the expression over time of the known and the proposed novel LSEC markers (as described in **Figure 3F**). RT-qPCR demonstrated that the LSEC markers *FCGR2B* (p<0.001), *LYVE1* (p<0.001), *MRC1* (p=0.002), *CRHBP* (p=0.025), *FCN3* (p=0.011), and *OIT3* (p=0.018) were significantly higher expressed in ETV2-SPI1-ECs compared to ETV2-ECs **(Figure 6C** and **Supplementary Figure 7C)**. This was further substantiated by immunostaining (**Supplementary Figure 8**) and flow cytometry analysis **(Figure 6E** and **Supplementary Figure 3)**, which showed a higher expression of CD32B and MRC1 in the ETV2-SPI1-ECs compared to the ETV2-ECs. CLEC4G, a highly specific LSEC marker, was however not expressed in either ETV2-ECs or ETV2-SPI1-ECs (data not shown).

As *ETV2* is only expressed until E10.5 in the mouse (53), we tested if persistent induction of *ETV2* might prohibit differentiation of ETV2-ECs into ECs with a more mature, LSEC-like phenotype. Therefore, to replace *ETV2* overexpression with TFs that are expressed in adult ECs, we cotransduced lentiviral vectors encoding the *SPI1, FLI1*, and/or *ERG* genes in ETV2-ECs on day 6 of differentiation, followed by doxycycline omission, and hence loss of *ETV2* expression, as of day 8. Loss of ETV2 expression caused a significant drop (>90%, p<0.01) in *PECAM1* expression unless cells were cotransduced with FLI1 and/or ERG **(Supplementary Figure 2A)**. Combined induction of *ETV2, ERG*, and *SPI1* led to increased expression of *MRC1* and *OIT3* (p<0.05). In contrast, combined induction of *FLI1* and *SPI1* decreased expression of *FCN3* and *DNASE1L3* (p<0.05) **(Supplementary Figure 2B-C)**. Overall, replacement of *ETV2* overexpression with *FLI1* or *ERG* overexpression could maintain the endothelial phenotype, but did not further enhance LSEC marker expression.

As the differentiation assay did not include VEGFA, and VEGFA may be beneficial for the LSEC phenotype (17,54), we repeated differentiations with inclusion of VEGFA from day 6 onwards. Expression of *LYVE1* in ETV2-SPI1-ECs was significantly induced by VEGFA (19.71-fold; p=0.005) **(Supplementary Figure 1)**. However, VEGFA did not significantly induce other LSEC markers such as *FCGR2B, STAB1*, and *STAB2*.

### Doxycycline-inducible *SPI1* overexpression improves LSEC features in ETV2-ECs

To assess LSEC functionality of ETV2-SPI1-ECs, we performed assays for the uptake of formaldehyde-treated FITC-coupled albumin (FSA-FITC) **(Figure 7A-C)** as well as IgG-AF555 **(Figure 7D-F)**. FSA-FITC uptake, which is scavenger receptor-mediated, is efficient in primary mouse LSEC **(Figure 7B)**. In PSC-derived ETV2-ECs, FSA-FITC uptake was enhanced in ETV2-SPI1-ECs when compared with ETV2-ECs (p<0.001). Additionally, when VEGFA was supplemented to the culture media, the uptake of FSA-FITC was increased in both cell populations (p<0.001) **(Figure 7C)**. Furthermore, CD32B-mediated IgG uptake was observed in only 22.21% (301/1355) of ETV2-ECs **(Figure 7D-E)**, while we could detect IgG-AF555 uptake in 68.60% (933/1360) of ETV2-SPI1-ECs (p=0.022), with the IgG-AF555 fluorochrome co-localised with endosomes identified by a RAB5 antibody (Pearson correlation for co-localisation: 0.733 ± 0.055) **(Figure 7G)**. Additionally, the IgG-AF555-positive ETV2-SPI1-ECs were significantly brighter than the IgG-AF555-positive ETV2-ECs (p<0.001) **(Figure 7F)**.

We also performed scanning electron microscopy on ETV2-ECs and ETV2-SPI1-ECs (with mouse LSECs as control) to determine if induction of *SPI1* induced fenestrae formation **(Figure 7H)**. Mouse LSECs showed clear fenestrations grouped in sieve plates. LSEC fenestration diameter was 122.73 ± 48.26 nm **(Figure 7I)**. However, neither ETV2- or ETV2-SPI1-ECs displayed fenestrations. Some transcytoplasmic holes were observed in ETV2-ECs and ETV2-SPI1-ECs but these holes were much smaller, i.e. 28.38 ± 18.02 nm and 33.55 ± 22.79 nm respectively, and likely due to SEM preparation artefacts as previously indicated by Elvevold et al. (2007) (55). We did note a clear morphological difference; ETV2-SPI1-ECs appeared more rounded with a few large cellular protrusions radiating from the main cell body, while ETV2-ECs appeared as flat cells with no or small and thin protrusions.

## Discussion

To identify TFs that are crucial for LSEC fating, we created a computational workflow (*CenTFinder)* that applies several bioinformatic tools on (a meta-analysis of) transcriptomics datasets. Our analysis identified *SPI1* as a key TF in a number of immune response pathways characteristic for LSECs. We demonstrated that forced expression of *SPI1* in PSC-derived ECs can partially fate PSC-ECs to an LSEC-like phenotype.

Classically, in differential expression analyses genes or TFs are ranked by fold change and/or p-value. However, such ranking may not necessarily yield (all) relevant TFs, as is exemplified by *SPI1*, which did not rank in the top 20 of differentially expressed TFs in the *CenTFinder* analysis. TFs might be differentially expressed due to differences between *in vivo* and *in vitro* environments or due to impurities in primary cell populations. Additionally, the fold change of a given TF does not provide knowledge regarding its number of dowstream targets, regarding its relevance to the cell-of-interest’s phenotype, or regarding the presence and complexity of its interactions with other proteins or signalling molecules.

To address these issues and to identify TFs that are crucial for LSEC fating, *CenTFinder* combines gene co-expression analysis (WGCNA), TF binding motif enrichment analysis (RcisTarget), and differential gene expression analysis. Several studies have been published that combine microarray meta-analysis with gene co-expression methods (56,57), however to date only a single study has combined this with TF binding motif enrichment as well (58). Furthermore, the SCENIC workflow (39), a well-established analysis tool for single-cell RNA sequencing data, is in essence based on similar premises.

The underlying rationale for gene co-expression analyses is that genes that are associated with the same biological process or pathway tend to be coregulated, thereby increasing the correlation between their expression patterns. Expression coregulation implies the existence of a subset of responsible upstream TFs, which hence would display higher connectivity and cluster-centrality (kME). Applying TF binding motif enrichment analysis on gene co-expression modules instead of the full gene set decreased the likelihood of prioritising a TF that was assigned to a module due to mere spurious correlation (e.g. correlation due to array probe position (59)). Simultaneously, it increased the likelihood of prioritising relevant TFs, as these TFs would not only be highly co-expressed with their module members but would also have enriched binding motifs in the cis-regulatory elements of these very same module members. Furthermore, TFs with enriched binding motifs in relatively smaller clusters (such as *SPI1* in our analysis) might not be detected by binding motif enrichment applied on the full list of differentially expressed genes. TFs were constrained by their centrality in a functional co-expression module and by the presence of enriched binding motifs. As TFs are assigned to a functional co-expression module, one can disregard TFs of modules that are not specifically related to the cell type of interest. For instance, in our analysis we found that only three modules (immune response) out of seven were relevant for the LSEC phenotype and hence we excluded TFs from other modules. Finally, we prioritised TFs that were differentially expressed between ETV2-ECs and LSECs as we reasoned that correcting the respective module activities might suffice by mere overexpression of these TFs.

We note that such strategy could be of interest as well in any other research aiming at improved differentiation and specification of PSC-progeny towards a cell type of interest. Additionally, the developed package can also identify TFs central to response to chemicals or different culturing conditions. The only limiting step in the described strategy is the number of arrays required for gene co-expression inference. As per their FAQ page (https://horvath.genetics.ucla.edu/html/CoexpressionNetwork/Rpackages/WGCNA/faq.html), the WGCNA authors do not recommend applying WGCNA on less than 20 samples. However, to counter noise, false-positive associations, and hence incorrect TF binding site enrichments, we would use at least 100 arrays. Extensive tutorials have been made available for WGCNA and RcisTarget analysis by their respective authors. Similarly, we made a tutorial available at https://github.com/jonathandesmedt92/CenTFinder.

We identified seven WGCNA modules, of which six were differentially active in LSECs compared to other ECs. Consistent with the known roles of LSECs in innate and immune functions, three modules contained genes related to immune response pathways. Interestingly, *SPI1* was identified as one of the key regulators of two of these immune clusters. *SPI1*, the gene encoding for the PU.1 protein, is commonly believed to play a pivotal role in B cells, NK cells, granulocytes, dendritic cells and macrophages (60–63). Although PU.1 and IRF TFs were recently identified as TFs that are potentially relevant for the LSEC phenotype (64), to our knowledge no group has functionally validated its role in fating ECs to LSEC-like cells.

We here provide conclusive evidence for PU.1 to be a master regulator for LSEC fating and as a key regulatory role in LSEC functionality. Over a hundred transciptional targets for PU.1 in haematopoietic progeny had already been identified by various groups (65), including *CLEC4G, FCGR2B*, and *MRC1*. Although *CLEC4G* expression was not induced upon *SPI1* overexpression in PSC-ECs, other established LSEC markers, such as *FCGR2B* and *MRC1*, were significantly induced. Furthermore, novel putative LSEC markers identified in this study (and in others (42,45,50)) such as *FCN2, FCN3, CRHBP*, and *OIT3* were also induced by PU.1. Overexpression of PU.1 induced not only increased expression of *FCGR2B*, but the activity of the CD32B receptor was also significantly higher, as was evidenced by IgG-AF555 uptake. Finally, uptake of FSA-FITC, a commonly used functional assay for LSECs, was more evident in ETV2-SPI1-ECs but not in ETV2-ECs, and this effect was enhanced with VEGFA supplementation, which is known to support *in vivo* LSEC differentiation (66,67) as well as *in vitro* culture (17). However, as STAB1/2 expression remained unchanged upon VEGFA addition, the mechanism could be post-translational, for instance at the level of receptor recycling.

While PU.1 is already known to regulate expression of cytokines, antibodies, and antibody receptors in a variety of cell types, to date no group, to our knowledge, described this in LSECs. Even more, it has been described that LSECs express CD45, a common haematopoietic marker (68). Considering this, the fact that PU.1 is a critical TF for haemotopoietic stem cells, and given the unique immune roles of LSECs, it raises the question whether the LSEC might descend from an intermediate cell type in the haemogenic endothelium. Future lineage tracing experiments during development will be required to help define LSEC identity and origin. Based on the very recent single-cell RNAseq data of Wang et al. 2020 (69) on human developing liver, it might also be of interest to test if specific TFs that are for instance present in endocardium during development, may aid in generating LSECs from PSCs.

Further research could also address the role of other TFs that are central in the immune response modules. Moreover, TFs that appear cluster-central and have enriched binding motifs within their respective modules but did not pass stringent differential expression criteria, could also be evaluated. The activity (or mere nuclear translocation) of such TFs could be modulated by interacting small molecules or other signalling factors. Furthermore, modules with lower activity in LSECs might contain TFs that interfere with immune response modules or TFs inhibiting the formation of fenestrae. Hence, knockdown of such TFs might further improve the LSEC phenotype.

In summary, our findings combine complementary transcriptomics analysis methods in order to maximise knowledge extraction from transcriptome data sets. For this purpose we developed the *CenTFinder* R package which bundles meta-analysis, gene co-expression analysis, TF binding motif enrichment analysis, and differential expression analysis in one integrated pipeline, providing also a framework for identifying TFs involved in, for instance, other endothelial lineage fating of stem cells. Based on this approach, we identified *SPI1* as the most highly ranked LSEC-specific TF, and demonstrated the importance of PU.1 in LSEC differentiation from pluripotent stem cells.

## Acknowledgements

We would like to thank Prof.Dr. Stein Aerts for his valuable input, advice, and discussions on bioinformatic strategies, Dr. Sara Aibar for her advice on the usage of RcisTarget, Dr. Cristina Øie for her critical assessment of and advice on the IgG uptake assays, and Dr. Aernout Luttun for his well-valued advice. Furthermore, we would like to acknowledge the VIB Nucleomics Core for the RNA sequencing, related quality control, and fruitful discussions. Confocal Images were recorded on a Zeiss LSM 880 – Airyscan (Cell and Tissue Imaging Cluster (CIC), Supported by Hercules AKUL/15/37_GOH1816N and FWO G.0929.15 to Pieter Vanden Berghe, University of Leuven. This study was funded by grants to J.D.S (Fonds Wetenschappelijk Onderzoek 1S33916N), R.B (Agentschap voor Innovatie door Wetenschap en Technologie SB-121393), IWT-140045, HILIM-3D to C.M.V. and L.v.G, FWO-G0D9917N to C.V., and funding from the EU-ToxRisk, a project running under the European Union’s Horizon 2020 research and innovation programme, grant agreement No 681002.

## Author contributions

J.D.S wrote all code for analysis and developed the *CenTFinder* package. J.D.S and I.T. analysed and interpreted the *CenTFinder* analysis and performed the lentiviral transductions. J.D.S. and S.G. performed the RT-qPCR, immunostainings, and cell culture. J.D.S., S.Z., B.T., S.G., and S.V. all contributed to (the set-up of) the flow cytometry readings. Confocal imaging and colocalisation analysis was entirely performed by B.T.. M.A.V. and R.B. generated the H9-ETV2 stem cell line and performed initial characterisations. J.D.S. generated the H9-ETV2-SPI1 stem cell line. R.F.M.D.C. performed the computational image analysis on immunostainings and the IgG uptake assays. J.D.S. and P.B. performed the SEM preparation and P.B. did the SEM imaging. Primary mouse cells were isolated by E.A.v.O.. FSA-FITC uptake assays were performed by E.A.v.O. and A.S. and IgG-AF555 uptake assays were performed by J.D.S.. All statistical analyses were performed by J.D.S.. J.D.S, L.A.v.G., S.A., and C.V. conceived the main concept of the presented study. J.D.S. and C.V. wrote majority of the manuscript. All authors contributed to the final version of the manuscript, as well as to discussions about and interpretations of the presented results.

