## Supplementary Figures for "PU.1 drives specification of pluripotent stem cell-derived endothelial cells to LSEC-like cells"

### Slide 1
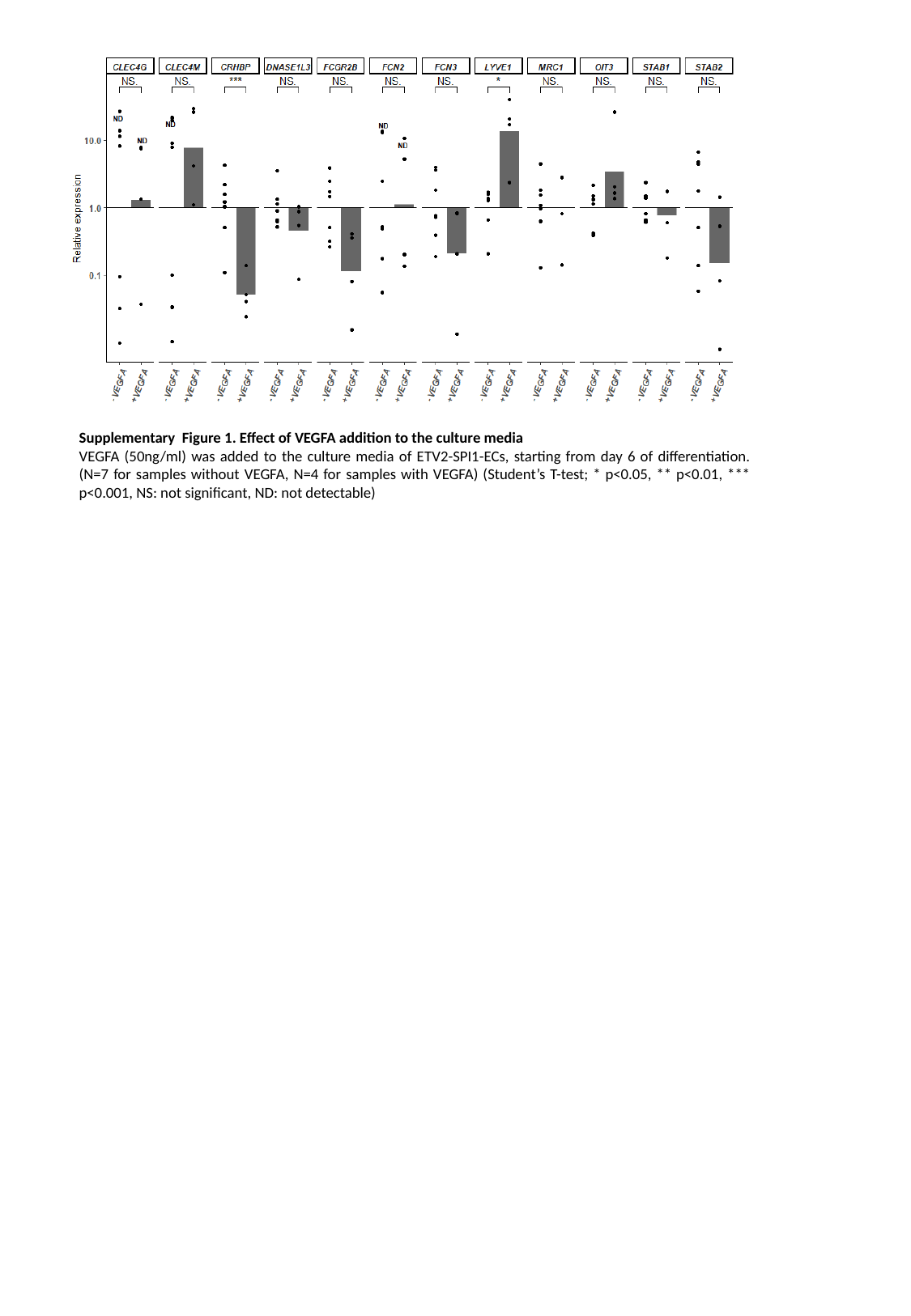

Supplementary Figure 1. Effect of VEGFA addition to the culture media
VEGFA (50ng/ml) was added to the culture media of ETV2-SPI1-ECs, starting from day 6 of differentiation. (N=7 for samples without VEGFA, N=4 for samples with VEGFA) (Student’s T-test; * p<0.05, ** p<0.01, *** p<0.001, NS: not significant, ND: not detectable)

### Slide 2
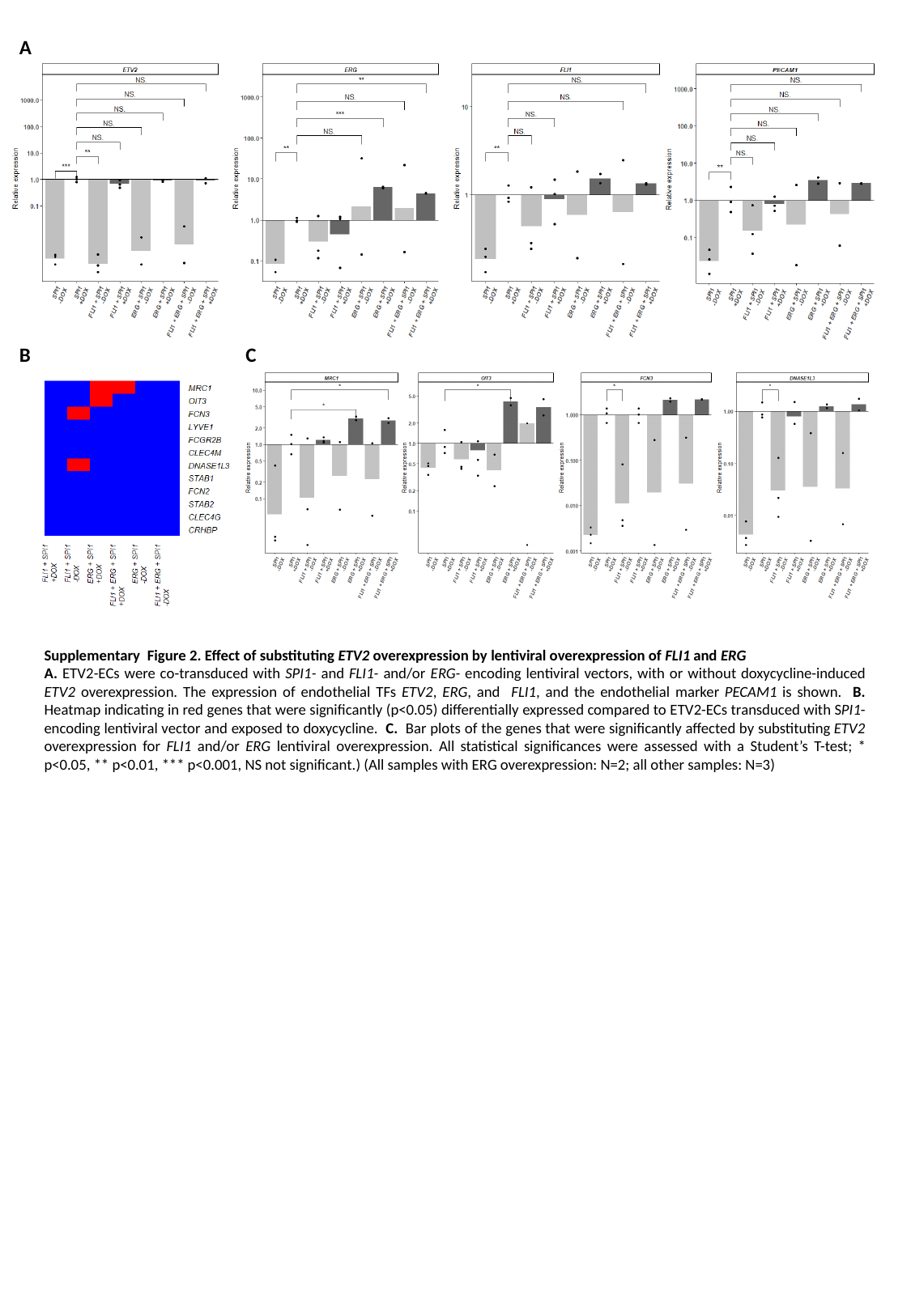

A
B
C
Supplementary Figure 2. Effect of substituting ETV2 overexpression by lentiviral overexpression of FLI1 and ERG
A. ETV2-ECs were co-transduced with SPI1- and FLI1- and/or ERG- encoding lentiviral vectors, with or without doxycycline-induced ETV2 overexpression. The expression of endothelial TFs ETV2, ERG, and FLI1, and the endothelial marker PECAM1 is shown. B. Heatmap indicating in red genes that were significantly (p<0.05) differentially expressed compared to ETV2-ECs transduced with SPI1-encoding lentiviral vector and exposed to doxycycline. C. Bar plots of the genes that were significantly affected by substituting ETV2 overexpression for FLI1 and/or ERG lentiviral overexpression. All statistical significances were assessed with a Student’s T-test; * p<0.05, ** p<0.01, *** p<0.001, NS not significant.) (All samples with ERG overexpression: N=2; all other samples: N=3)

### Slide 3
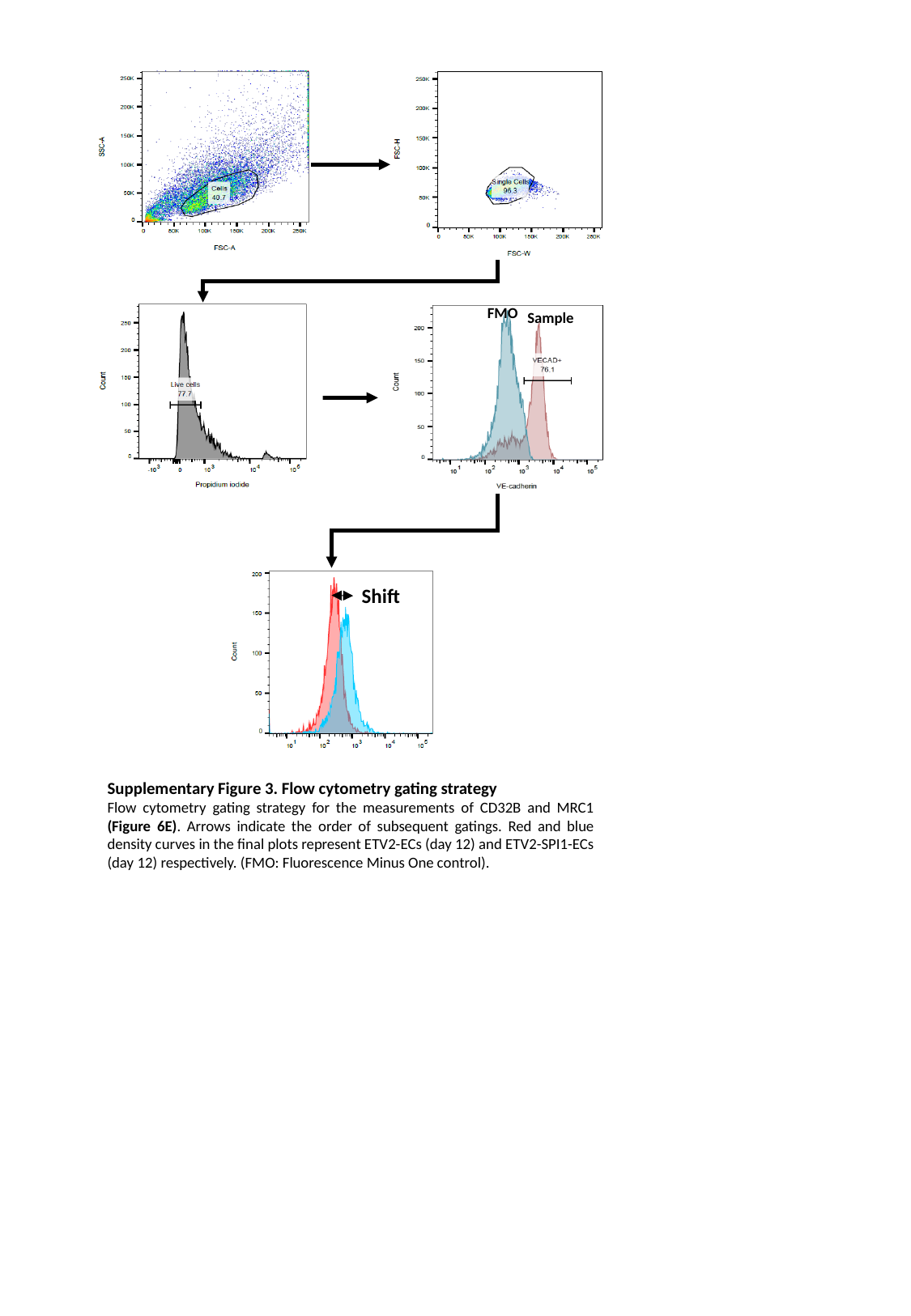

FMO
Sample
Shift
Supplementary Figure 3. Flow cytometry gating strategy
Flow cytometry gating strategy for the measurements of CD32B and MRC1 (Figure 6E). Arrows indicate the order of subsequent gatings. Red and blue density curves in the final plots represent ETV2-ECs (day 12) and ETV2-SPI1-ECs (day 12) respectively. (FMO: Fluorescence Minus One control).

### Slide 4
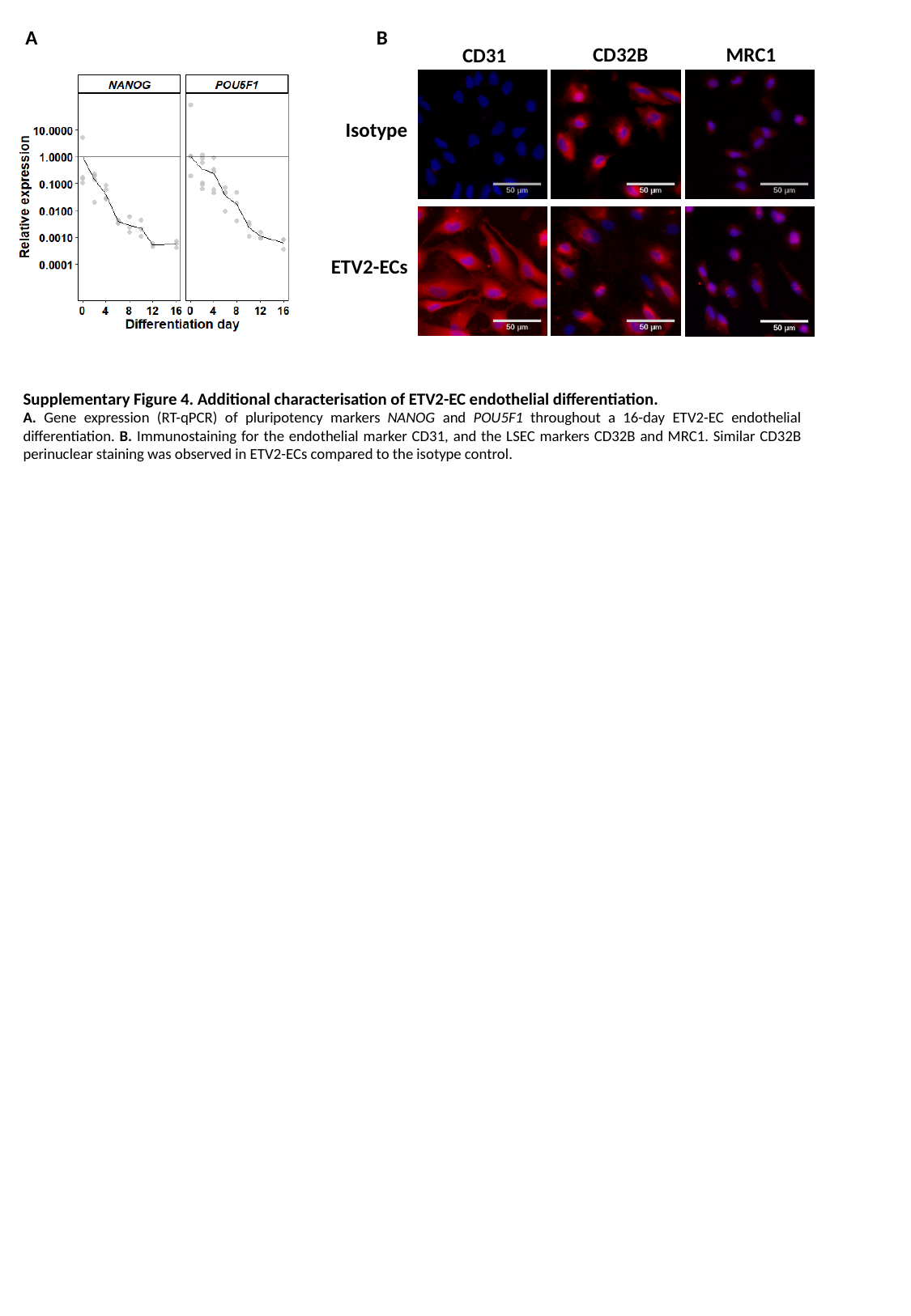

B
A
MRC1
CD32B
CD31
Isotype
ETV2-ECs
Supplementary Figure 4. Additional characterisation of ETV2-EC endothelial differentiation.
A. Gene expression (RT-qPCR) of pluripotency markers NANOG and POU5F1 throughout a 16-day ETV2-EC endothelial differentiation. B. Immunostaining for the endothelial marker CD31, and the LSEC markers CD32B and MRC1. Similar CD32B perinuclear staining was observed in ETV2-ECs compared to the isotype control.

### Slide 5
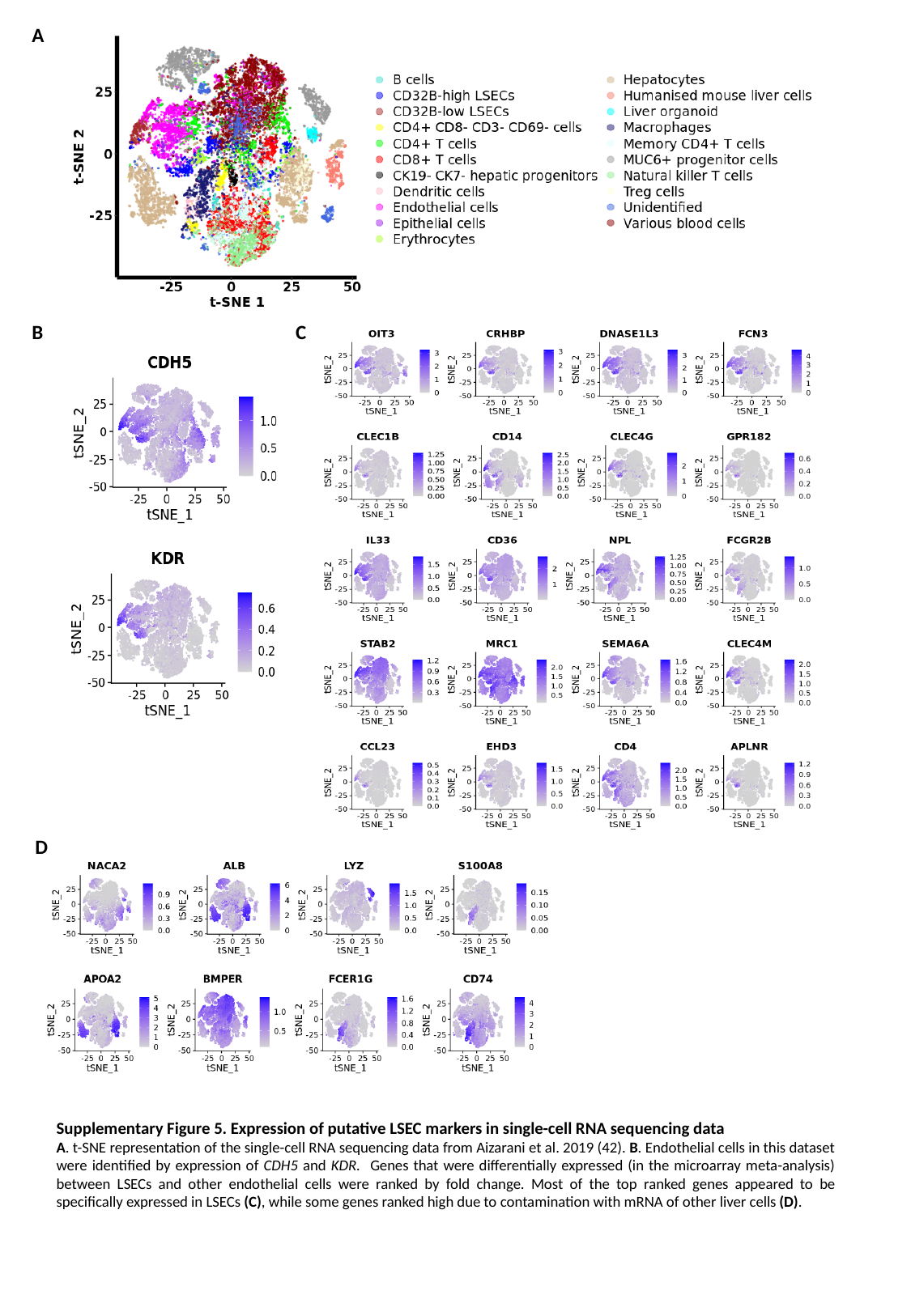

A
C
B
D
Supplementary Figure 5. Expression of putative LSEC markers in single-cell RNA sequencing data
A. t-SNE representation of the single-cell RNA sequencing data from Aizarani et al. 2019 (42). B. Endothelial cells in this dataset were identified by expression of CDH5 and KDR. Genes that were differentially expressed (in the microarray meta-analysis) between LSECs and other endothelial cells were ranked by fold change. Most of the top ranked genes appeared to be specifically expressed in LSECs (C), while some genes ranked high due to contamination with mRNA of other liver cells (D).

### Slide 6
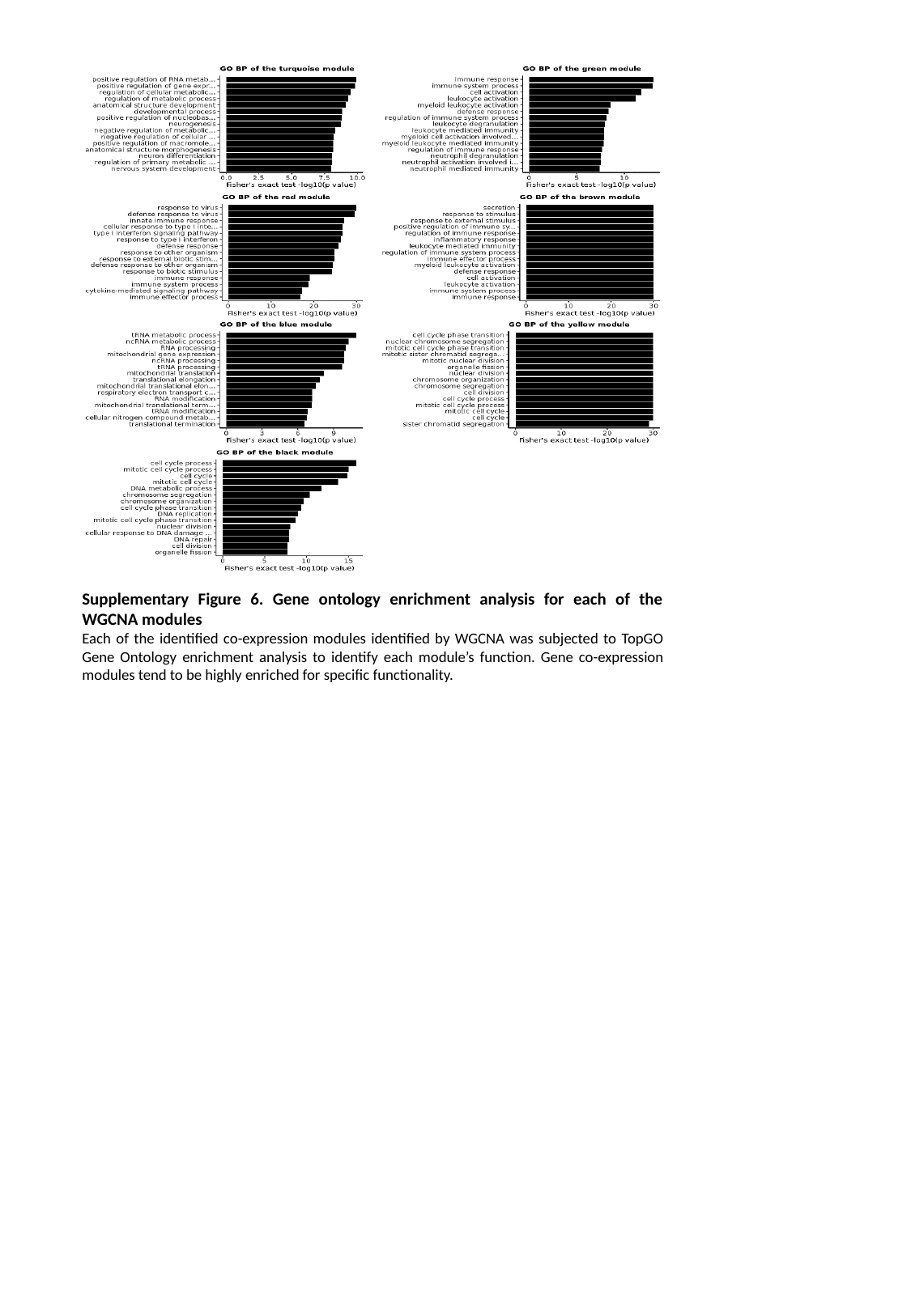

Supplementary Figure 6. Gene ontology enrichment analysis for each of the WGCNA modules
Each of the identified co-expression modules identified by WGCNA was subjected to TopGO Gene Ontology enrichment analysis to identify each module’s function. Gene co-expression modules tend to be highly enriched for specific functionality.

### Slide 7
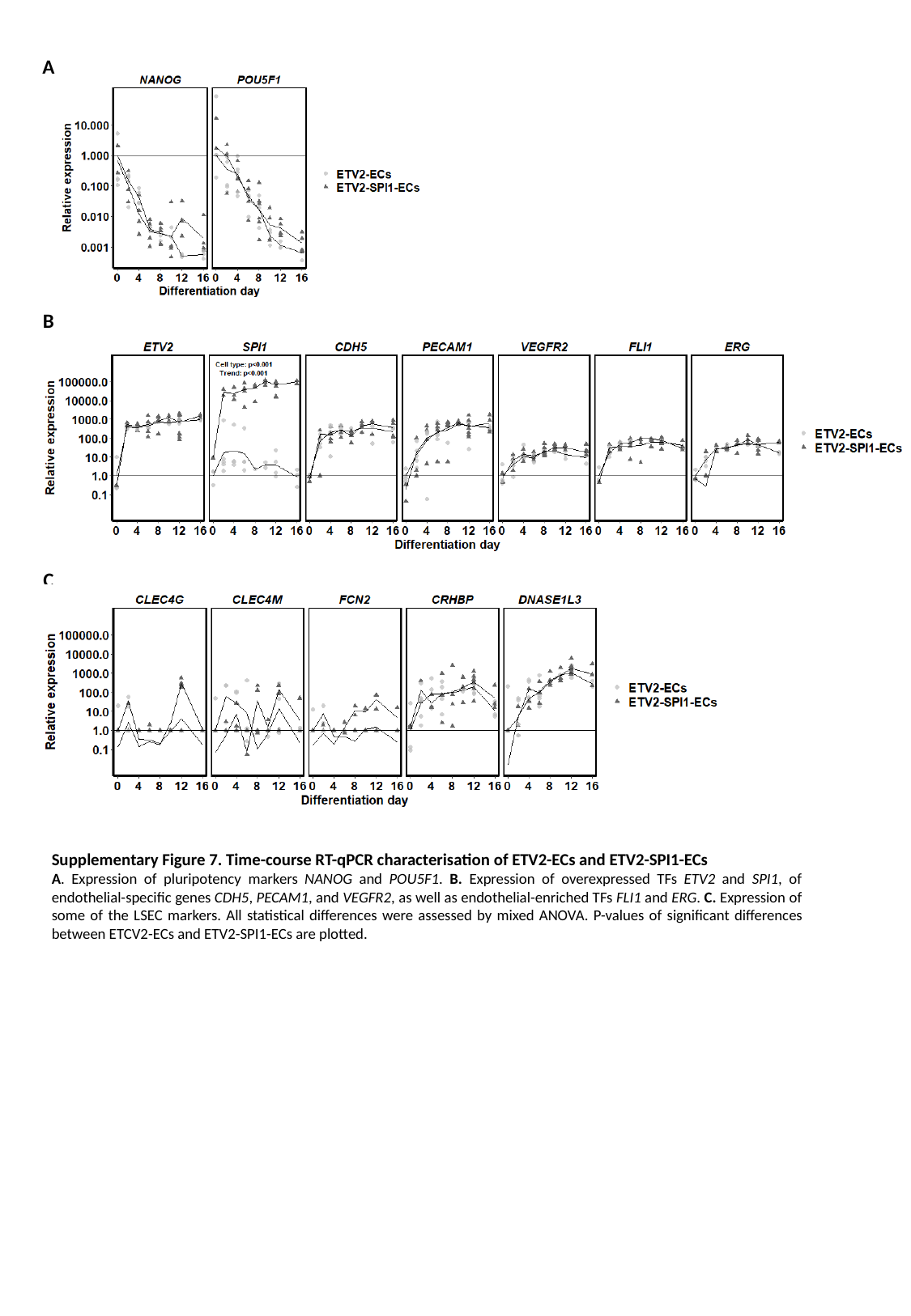

A
B
C
Supplementary Figure 7. Time-course RT-qPCR characterisation of ETV2-ECs and ETV2-SPI1-ECs
A. Expression of pluripotency markers NANOG and POU5F1. B. Expression of overexpressed TFs ETV2 and SPI1, of endothelial-specific genes CDH5, PECAM1, and VEGFR2, as well as endothelial-enriched TFs FLI1 and ERG. C. Expression of some of the LSEC markers. All statistical differences were assessed by mixed ANOVA. P-values of significant differences between ETCV2-ECs and ETV2-SPI1-ECs are plotted.

### Slide 8
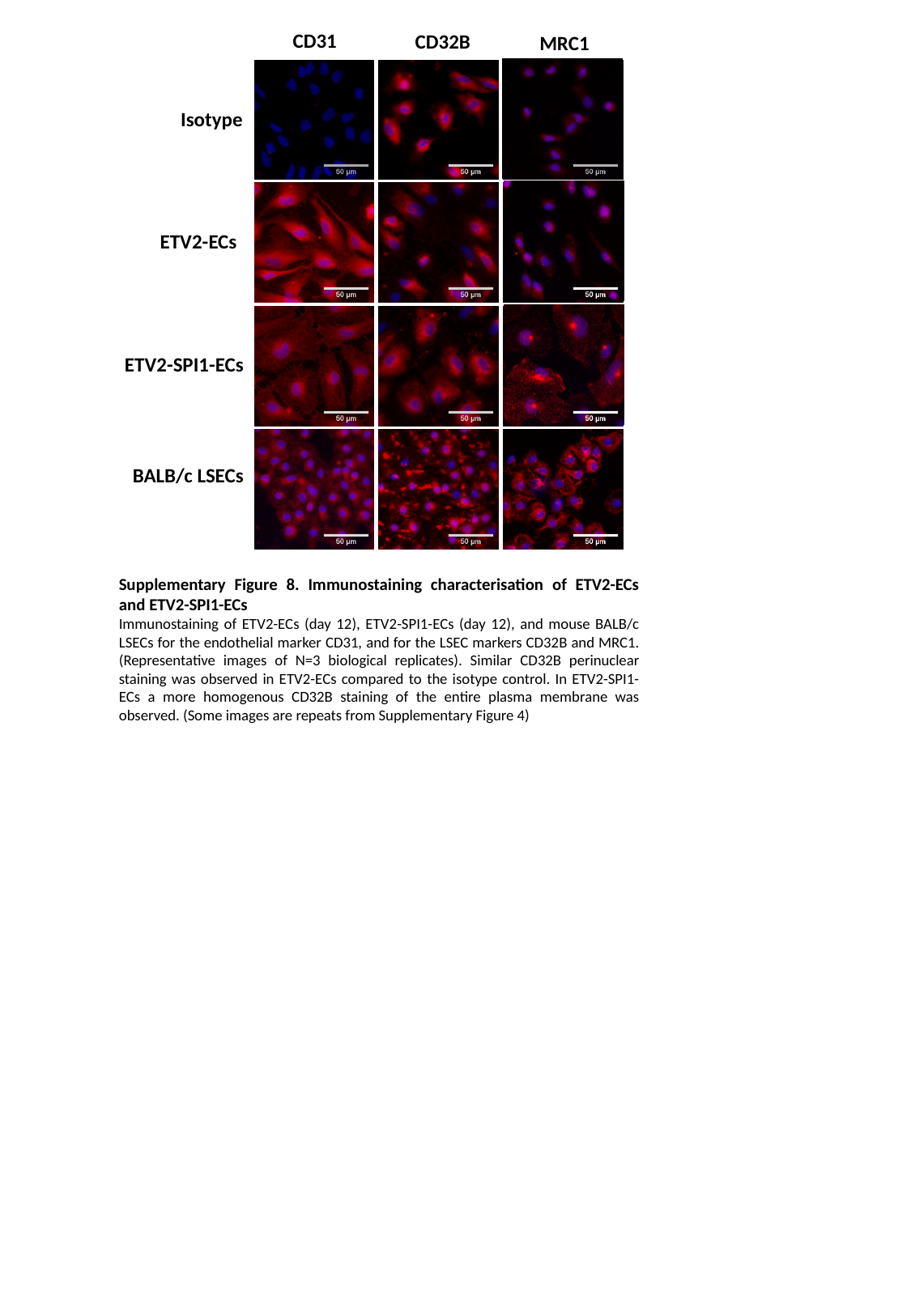

CD31
CD32B
MRC1
Isotype
ETV2-ECs
ETV2-SPI1-ECs
BALB/c LSECs
Supplementary Figure 8. Immunostaining characterisation of ETV2-ECs and ETV2-SPI1-ECs
Immunostaining of ETV2-ECs (day 12), ETV2-SPI1-ECs (day 12), and mouse BALB/c LSECs for the endothelial marker CD31, and for the LSEC markers CD32B and MRC1. (Representative images of N=3 biological replicates). Similar CD32B perinuclear staining was observed in ETV2-ECs compared to the isotype control. In ETV2-SPI1-ECs a more homogenous CD32B staining of the entire plasma membrane was observed. (Some images are repeats from Supplementary Figure 4)
