## Supplementary Tables for "PU.1 drives specification of pluripotent stem cell-derived endothelial cells to LSEC-like cells"

**Supplementary Table 1 RT-qPCR primer sequences**

| **Intended target** | **Gene** | **Forward primer** | **Reverse primer** |
| --- | --- | --- | --- |
| Stem cells | *NANOG* | CCTGTGATTTGTGGGCCTG | GACAGTCTCCGTGTGAGGCAT |
|  | *POU5F1* | TCGAGAAGGATGTGGTCCGA | GCCTCAAAATCCTCTCGTTG |
| Endothelial cells | *CDH5* | GTTCACGCATCGGTTGTTC | TCTGCATCCACTGCTGTCA |
|  | *ERG* | TCCTCCAGCGACTATGGACA | TTTGATGGTGACCCTGGCTG |
|  | *ETV2* | CATCCGTTGGACTGGCAACA | TTCATGCCCGGCTTTCTCTT |
|  | *FLI1* | GAAGAGGAGCTTGGGGCAAT | GGGCCGTTGCTCTGTATTCT |
|  | *PECAM1* | TCTGCACTG CAGGTATTGACAA | CTGATCGATTCGCAACGGA |
|  | *VEGFR2* | ACAACCAGACGGACAGTGGT | AGCCTTCAGATGCCACAGAC |
| LSECs | *CLEC4G* | CTTCCTCACTCGGAACACG | GGTCAGCAGTTGTGCCTTTT |
|  | *CLEC4M* | CTGTCCCAAGGACTGGACAT | CGTGCCTTCCTGATTTAGGT |
|  | *CRHBP* | GGGAGGAACTGGATTGGACC | CAACTTTCATCTGGGCCGGG |
|  | *DNASE1L3* | CGCTGGAAGGCGGAGAATTT | CCTTCTTGGGGACGTAGCTG |
|  | *FCGR2B* | CAAAGTTGGGGCTGAGAACA | CCCTGTCCTCCCCAAGGGGAA |
|  | *FCN2* | GGGCAGTGCGGGAGATTC | CACAGCACAATTTCCGGTGTT |
|  | *FCN3* | CTTACTCTCCAGGGTAACTGGG | GGGCGAAAGTACGGTTACCA |
|  | *LYVE1* | TTTGCAGCCTATTGTTACAACTCAT | GGGATGCCACCCAGTAGGTA |
|  | *MRC1* | TCCTGTCCATCAGGAGAAGG | ATTTCTGTGATTCGGCATCC |
|  | *OIT3* | TGGTACAGTGGTCGATGTGG | GGTCTGCTTGGGTAGACCTG |
|  | *SPI1* | GGAGAGCCATAGCGACCATT | CTGGAGCTCCGTGAAGTTGT |
|  | *STAB1* | TCTACCGATCTGGCTTCTCC | CCAGCTCACACACCTCACAG |
|  | *STAB2* | GCAAGAAGATGTGATAGGAAGTCTC | ACAACACCGAGGTTGGAGAT |
| Reference gene | *EEF1A1* | CAGAACACAGGTGTCGTGAA | CCGAATCTACGTGTCCAATGA |
|  | *GAPDH* | TCAAGAAGGTGGTGAAGCAGG | ACCAGGAAATGAGCTTGACAAA |
|  | *RPL19* | ATTGGTCTCATTGGGGTCTAAC | AGTATGCTCAGGCTTCAGAAGA |
|  | *RPS23* | GGTTGCTTGAACTTTATTGAGGAAA | AGCGGACTCCAGGAATATCA |

**Supplementary Table 2 Antibodies and dyes**

| **ID** | **Antibody/Dye** | **Clonality** | **Catalogue number** | **Company** | **Fluorophore** | **Application** |
| --- | --- | --- | --- | --- | --- | --- |
| 1 | CD144 (VE-cadherin) Monoclonal Antibody | Monoclonal (16B1) | 13-1449-82 | ThermoFisher Scientific | Biotin – ID 12 was used as secondary dye | Flow cytometry |
| 2 | Anti-VE Cadherin antibody - Intercellular Junction Marker | Polyclonal | ab33168 | Abcam | Unconjugated – ID 11 was used as secondary antibody | Immunostaining |
| 3 | Anti-LYVE1 antibody | Polyclonal | ab14917 | Abcam | Unconjugated – ID 11 and 10 were used as secondary antibody for staining and FC respectively | Immunostaining Flow cytometry |
| 4 | Anti-Mannose Receptor antibody | Polyclonal | ab64693 | Abcam | Unconjugated – ID 11 and 10 were used as secondary antibody for staining and FC respectively | Immunostaining Flow cytometry |
| 5 | CD32b antibody | Polyclonal | orb44658 | Biorbyt | Unconjugated – ID 11 was used as secondary antibody | Immunostaining |
| 6 | Anti-CD32B antibody – C-terminal | Polyclonal | ab151497 | Abcam | Unconjugated – ID 10 was used as secondary antibody | Flow cytometry |
| 7 | Human LSECtin/CLEC4G Alexa Fluor 647-conjugated Antibody | Monoclonal (845404) | FAB2947R-100UG | R&D Systems | Conjugated with Alexa Fluor 647 | Flow cytometry |
| 8 | Anti-CD45 antibody | Polyclonal | ab10558 | Abcam | Unconjugated – ID 10 was used as secondary antibody | Flow cytometry |
| 9 | Rab5 (C8B1) Rabbit mAb #3547 | Monoclonal (C8B1) | #3547 | Cell Signaling Technology | Unconjugated – ID 10 was used as secondary antibody | Immunostaining |
| 10 | Donkey anti-Rabbit IgG (H+L) Highly Cross-Adsorbed Secondary Antibody, Alexa Fluor 647 | Polyclonal | A-31573 | ThermoFisher Scientific | Conjugated with Alexa Fluor 647 | Immunostaining  Flow cytometry |
| 11 | Donkey anti-Rabbit IgG (H+L) Highly Cross-Adsorbed Secondary Antibody, Alexa Fluor 555 | Polyclonal | A-31572 | ThermoFisher Scientific | Conjugated with Alexa Fluor 555 | Immunostaining  IgG uptake assay |
| 12 | eBioscience™ Streptavidin eFluor™ 450 Conjugate | NA | 48-4317-82 | ThermoFisher Scientific | Conjugated with eFluor™ 450 | Flow cytometry |
| 13 | Propidium iodide | NA | 81845-25MG (80513) | Sigma Aldrich | Inherent | Flow cytometry |
| 14 | Alexa Fluor™ 488 Phalloidin | NA | A12379 | ThermoFisher Scientific | Conjugated with Alexa Fluor 488 | Immunostaining |
| 15 | Prolong Gold antifade reagent with DAPI | NA | P-36931 | ThermoFisher Scientific | Inherent | Immunostaining |
| 16 | Donkey anti-Mouse IgG (H+L) Highly Cross-Adsorbed Secondary Antibody, Alexa Fluor 647 | Polyclonal | A-31571 | ThermoFisher Scientific | Conjugated with Alexa Fluor 647 | Immunostaining |

**Supplementary Table 3 Microarrays included in the CenTFinder analysis**

| **Cell type** | **File name** | **Platform** | **ArrayExpress accession number** |
| --- | --- | --- | --- |
| blood outgrowth EC | GSM952933_LDS1.CEL | HG-U133_Plus_2 | E-GEOD-38961 |
|  | GSM952934_BC248.CEL |  |  |
|  | GSM952935_LDS5_OEC.CEL |  |  |
|  | GSM952936_BC14_OEC.CEL |  |  |
|  | GSM952937_LDS11_OEC.CEL |  |  |
|  | GSM952938_BC401_OEC.CEL |  |  |
|  | NUID-0000-0097-6615.cel |  | E-MTAB-2495 |
|  | NUID-0000-0097-6618.cel |  |  |
|  | NUID-0000-0097-6624.cel |  |  |
|  | NUID-0000-0097-6639.cel |  |  |
|  | NUID-0000-0097-6643.cel |  |  |
|  | NUID-0000-0097-6646.cel |  |  |
|  | NUID-0000-0097-6649.cel |  |  |
|  | NUID-0000-0097-6665.cel |  |  |
|  | NUID-0000-0097-6666.cel |  |  |
|  | NUID-0000-0097-6670.cel |  |  |
|  | NUID-0000-0097-6674.cel |  |  |
|  | NUID-0000-0097-6689.cel |  |  |
|  | NUID-0000-0097-6692.cel |  |  |
|  | NUID-0000-0097-6693.cel |  |  |
|  | NUID-0000-0097-6696.cel |  |  |
| choroid EC | GSM524670.CEL | HG-U133_Plus_2 | E-MTAB-3732 |
| coronary artery EC | GSM1087598_S1_041311_SL_HGU133P2_041411.CEL | HG-U133_Plus_2 | E-GEOD-44596 |
|  | GSM1087599_S2_041311_SL_HGU133P2_041411.CEL |  |  |
|  | GSM1087600_S3_041311_SL_HGU133P2_041411.CEL |  |  |
|  | GSM1087601_C1_041311_SL_HGU133P2_041411.CEL |  |  |
|  | GSM1087602_C2_041311_SL_HGU133P2_041411.CEL |  |  |
|  | GSM1087603_C3_041311_SL_HGU133P2_041411.CEL |  |  |
|  | GSM1287224_RWX101025_Remaley_Vickers_HDL_AI1B.CEL | HuGene-1_0-st-v1 | E-GEOD-53201 |
|  | GSM1287225_RWX101025_Remaley_Vickers_HDL_AI2B.CEL |  |  |
|  | GSM1287226_RWX101025_Remaley_Vickers_HDL_AI3A.CEL |  |  |
|  | GSM1287227_RWX101025_Remaley_Vickers_HDL_C1.CEL |  |  |
|  | GSM1287228_RWX101025_Remaley_Vickers_HDL_C2.CEL |  |  |
|  | GSM1287229_RWX101025_Remaley_Vickers_HDL_C3.CEL |  |  |
|  | GSM1287230_RWX101025_Remaley_Vickers_HDL_DISCS1A.CEL |  |  |
|  | GSM1287231_RWX101025_Remaley_Vickers_HDL_DISCS2B.CEL |  |  |
|  | GSM1287232_RWX101025_Remaley_Vickers_HDL_DISCS3A.CEL |  |  |
|  | GSM1287233_RWX101025_Remaley_Vickers_HDL_DONOR1S1.CEL |  |  |
|  | GSM1287234_RWX101025_Remaley_Vickers_HDL_DONOR2S3.CEL |  |  |
|  | GSM1287235_RWX101025_Remaley_Vickers_HDL_DONOR3S3.CEL |  |  |
|  | GSM1287236_RWX101025_Remaley_Vickers_HDL_DONOR4S3.CEL |  |  |
|  | GSM1287237_RWX101025_Remaley_Vickers_HDL_DONOR5S2.CEL |  |  |
|  | GSM1287238_RWX101025_Remaley_Vickers_HDL_VES1.CEL |  |  |
|  | GSM1287239_RWX101025_Remaley_Vickers_HDL_VES2.CEL |  |  |
|  | GSM1287240_RWX101025_Remaley_Vickers_HDL_VES3.CEL |  |  |
|  | GSM1287241_RWX101025_Remaley_Vickers_HDL_WAI-1-1.CEL |  |  |
|  | GSM1287242_RWX101025_Remaley_Vickers_HDL_WAI-2-1.CEL |  |  |
|  | GSM1287243_RWX101025_Remaley_Vickers_HDL_WC1.CEL |  |  |
|  | GSM1287244_RWX101025_Remaley_Vickers_HDL_WC2.CEL |  |  |
|  | GSM1287245_RWX101025_Remaley_Vickers_HDL_WDISCS1-1.CEL |  |  |
|  | GSM1287246_RWX101025_Remaley_Vickers_HDL_WDISCS2-1.CEL |  |  |
|  | GSM1287247_RWX101025_Remaley_Vickers_HDL_WVES1.CEL |  |  |
|  | GSM1287248_RWX101025_Remaley_Vickers_HDL_WVES2.CEL |  |  |
| cultured aorta EC | GSM1063377_HAEC1_2.CEL | HG-U133_Plus_2 | E-GEOD-43475 |
|  | GSM1063378_HAEC2_9.CEL |  |  |
| cultured coronary artery EC | GSM1063379_HCAE_C1_11.CEL | HG-U133_Plus_2 | E-GEOD-43475 |
|  | GSM1063380_HCAEC2_9.CEL |  |  |
| cultured hepatic artery EC | GSM1063374_HA1_7.CEL | HG-U133_Plus_2 | E-GEOD-43475 |
|  | GSM1063375_HA2_1.CEL |  |  |
|  | GSM1063376_HA3_2.CEL |  |  |
| cultured hepatic vein EC | GSM1063409_HV1_4.CEL | HG-U133_Plus_2 | E-GEOD-43475 |
|  | GSM1063410_HV2_5.CEL |  |  |
|  | GSM1063411_HV3_6.CEL |  |  |
| cultured HUVEC | GSM1063403_HUVEC_C4_6.CEL | HG-U133_Plus_2 | E-GEOD-43475 |
| cultured iliac artery EC | GSM1063381_HIAE1_10.CEL | HG-U133_Plus_2 | E-GEOD-43475 |
|  | GSM1063382_HIAE4_8.CEL |  |  |
| cultured iliac vein EC | GSM1063383_HIVE1_5.CEL | HG-U133_Plus_2 | E-GEOD-43475 |
|  | GSM1063384_HIVE4_13.CEL |  |  |
|  | GSM1063385_HIVE5_14.CEL |  |  |
| cultured lung EC | GSM1185521_human_lungEC-control_1.CEL | HG-U133_Plus_2 | E-GEOD-48841 |
|  | GSM1185522_human_lungEC-control_2.CEL |  |  |
|  | GSM1185523_human_lungEC-DSCR-1_KD_1.CEL |  |  |
|  | GSM1185524_human_lungEC-DSCR-1_KD_2.CEL |  |  |
|  | GSM1185525_human_lungEC-control_VEGF0h.CEL |  |  |
| cultured lung EC | GSM1185526_human_lungEC-control_VEGF1h.CEL | HG-U133_Plus_2 | E-GEOD-48841 |
|  | GSM1185527_human_lungEC-control_VEGF_4h.CEL |  |  |
|  | GSM1185528_human_lungEC-control_VEGF_24h.CEL |  |  |
|  | GSM1185529_human_lungEC-Ad-DSCR1_VEGF0h.CEL |  |  |
|  | GSM1185530_human_lungEC-Ad-DSCR1_VEGF1h.CEL |  |  |
|  | GSM1185531_human_lungEC-Ad-DSCR1_VEGF4h.CEL |  |  |
|  | GSM1185532_human_lungEC-Ad-DSCR1_VEGF24h.CEL |  |  |
| cultured pulmonary artery EC | GSM1063386_HPAE1_1.CEL | HG-U133_Plus_2 | E-GEOD-43475 |
|  | GSM1063387_HPAE4_9.CEL |  |  |
|  | GSM1063388_HPAE5_1.CEL |  |  |
| cultured pulmonary vein EC | GSM1063389_HPVE2_4.CEL | HG-U133_Plus_2 | E-GEOD-43475 |
|  | GSM1063390_HPVE3_12.CEL |  |  |
| cultured umbilical artery EC | GSM1063391_HUAEC_C1_3.CEL | HG-U133_Plus_2 | E-GEOD-43475 |
|  | GSM1063392_HUAEC_C2_4.CEL |  |  |
|  | GSM1063393_HUAEC_C3_11.CEL |  |  |
|  | GSM1063394_HUAEC_C5_12.CEL |  |  |
|  | GSM1063395_HUAEC_C6_3.CEL |  |  |
| dermal microvascular EC | GSM617993.CEL | HG-U133_Plus_2 | E-MTAB-3732 |
|  | GSM617995.CEL |  |  |
|  | GSM617998.CEL |  |  |
|  | GSM618000.CEL |  |  |
|  | GSM618001.CEL |  |  |
|  | GSM618006.CEL |  |  |
| early endothelial precursor | GSM909291_S0119-139-UCB-eEPC.CEL | HG-U133_Plus_2 | E-GEOD-37045 |
| EC (unspecified) | GSM1624172_HUVEC-VGPCR.CEL | HG-U133_Plus_2 | E-GEOD-66503 |
|  | GSM1624173_HUVEC-PURO.CEL |  |  |
|  | GSM740042.CEL |  | E-GEOD-29881 |
|  | GSM740043.CEL |  |  |
|  | GSM740044.CEL |  |  |
|  | GSM740045.CEL |  |  |
|  | GSM740046.CEL |  |  |
|  | GSM740047.CEL |  |  |
|  | GSM740048.CEL |  |  |
|  | GSM740049.CEL |  |  |
|  | GSM740050.CEL |  |  |
|  | GSM740051.CEL |  |  |
|  | GSM740052.CEL |  |  |
|  | GSM740053.CEL |  |  |
|  | GSM740054.CEL |  |  |
|  | GSM740055.CEL |  |  |
|  | GSM740056.CEL |  |  |
|  | GSM740057.CEL |  |  |
|  | GSM740058.CEL |  |  |
|  | GSM740059.CEL |  |  |
|  | GSM740060.CEL |  |  |
|  | GSM740061.CEL |  |  |
| endothelial precursor | BEN_EPC.CEL | HuGene-1_0-st-v1 | E-MEXP-3071 |
|  | CUN-EPC.CEL |  |  |
|  | GSM663488.CEL | HG-U133_Plus_2 | E-GEOD-26950 |
|  | GSM663489.CEL |  |  |
|  | GSM663490.CEL |  |  |
|  | GSM663491.CEL |  |  |
|  | GSM663492.CEL |  |  |
|  | GSM663493.CEL |  |  |
|  | GSM663494.CEL |  |  |
|  | GSM663495.CEL |  |  |
|  | GSM663496.CEL |  |  |
|  | GSM663497.CEL |  |  |
|  | GSM663498.CEL |  |  |
| endothelial precursor | GSM663499.CEL | HG-U133_Plus_2 | E-GEOD-26950 |
|  | SHEN_EPC.CEL | HuGene-1_0-st-v1 | E-MEXP-3071 |
| fresh HUVEC | GSM1063407_HUVEC_F7_14.CEL | HG-U133_Plus_2 | E-GEOD-43475 |
|  | GSM1063408_HUVEC_F9_5.CEL |  |  |
| fresh umbilical artery EC | GSM1063396_HUAEC_F10_12.CEL | HG-U133_Plus_2 | E-GEOD-43475 |
|  | GSM1063397_HUAEC_F12_10.CEL |  |  |
|  | GSM1063398_HUAEC_F13_2.CEL |  |  |
|  | GSM1063399_HUAEC_F8_3.CEL |  |  |
| HUVEC | EXP1_1.4_W1.CEL | HuGene-1_0-st-v1 | E-MTAB-6521 |
|  | EXP1_CTRL_W3.CEL |  |  |
|  | EXP1_CTRL_W6.CEL |  |  |
|  | EXP2_CTRL_W3.CEL |  |  |
|  | EXP3_1.4_W1.CEL |  |  |
|  | EXP3_CTRL_W3.CEL | HuGene-1_0-st-v1 | E-MTAB-6521 |
|  | GSM1127001_NoRev2.CEL |  | E-GEOD-46248 |
|  | GSM1127004_Rev1.CEL |  |  |
|  | GSM1127007_Rev4.CEL |  |  |
| HUVEC | GSM1128073_EM_6_01292009_6_HOF.CEL |  | E-GEOD-46279 |
|  | HUVEC5_4h_FSAP_HuGene_1_0_st_v1.CEL | HuGene-1_0-st-v1 | E-MTAB-5592 |
|  | HUVEC8_10h_FSAP_HuGene_1_0_st_v1.CEL |  |  |
|  | HUVEC_USP10_siRNA_DLL4_2.CEL | HuGene-2_0-st | E-MTAB-7774 |
|  | MYC_siRNA_b.CEL |  | E-MTAB-4025 |
|  | GSM1099432_F1PS_05.CEL | HG-U133_Plus_2 | E-GEOD-45225 |
|  | GSM1099433_F1PS_06.CEL |  |  |
|  | GSM1099438_F5AS_11.CEL |  |  |
|  | GSM1184603_1_E4ORFCont1_UWS-KXX-120424-24-30585_1.CEL |  | E-GEOD-48786 |
|  | GSM1184604_2_E4ORFCont2_UWS-KXX-120424-24-30585_2.CEL |  |  |
|  | GSM1184607_8_E4ORFSort2_UWS-KXX-120424-24-30585_5.CEL |  |  |
|  | GSM1199152_101207_03_VEplusCsA1.CEL |  | E-GEOD-49426 |
|  | GSM1199154_101207_05_NFAT24.CEL |  |  |
|  | GSM458347.CEL |  | E-MTAB-3732 |
|  | GSM468615.CEL |  | E-GEOD-18913 |
|  | GSM468623.CEL |  |  |
|  | GSM476786.CEL |  | E-MTAB-3732 |
|  | GSM990632_MS517_I.1_101005.CEL |  | E-GEOD-40281 |
|  | GSM990633_MS517_Probe_IV_1_270905.CEL |  |  |
|  | GSM990635_MS517_II.2_101005.CEL |  |  |
|  | GSM990639_MS517_Probe_IV_4_270905.CEL |  |  |
| immortalised microvascular EC | GSM1035591_1_TIME_A.CEL | HG-U133_Plus_2 | E-GEOD-42216 |
|  | GSM1035592_2_TIME_B.CEL |  |  |
|  | GSM1035593_3_TIME_C.CEL |  |  |
|  | GSM1035594_4_TIME_D.CEL |  |  |
|  | GSM1035595_5_HMEC-1_A.CEL |  |  |
|  | GSM1035596_6_HMEC-1_B.CEL |  |  |
|  | GSM1035597_7_HMEC-1_C.CEL |  |  |
|  | GSM1035598_8_HMEC-1_D.CEL |  |  |
| intestinal lymphatic EC | GSM1466659_stat_siAS_B1.CEL | HuGene-1_0-st-v1 | E-GEOD-60152 |
|  | GSM1466660_stat_siAS_C7.CEL |  |  |
|  | GSM1466661_stat_siFoxC2_B2.CEL |  |  |
|  | GSM1466662_stat_siFoxC2_C8.CEL |  |  |
|  | GSM1466663_osc_siAS_B3.CEL |  |  |
|  | GSM1466664_osc_siAS_C9.CEL |  |  |
|  | GSM1466665_osc_siFoxC2_B4.CEL |  |  |
|  | GSM1466666_osc_siFoxC2_C10.CEL |  |  |
| iPSC EC | GSM1920951_CiRA00004_EC_HuGene-1_0-st-v1_.CEL | HuGene-1_0-st-v1 | E-GEOD-74453 |
|  | GSM1920952_CiRA00005_EC_HuGene-1_0-st-v1_.CEL |  |  |
|  | GSM1920953_CiRA00006_EC__HuGene-1_0-st-v1_.v2.CEL |  |  |
|  | GSM1920954_CiRA00007_EC_HuGene-1_0-st-v1_.CEL |  |  |
|  | GSM1920955_CiRA00008_EC_HuGene-1_0-st-v1_.CEL |  |  |
|  | GSM1920956_CiRA00009_EC_HuGene-1_0-st-v1_.CEL |  |  |
|  | GSM1920957_CiRA00010_EC_HuGene-1_0-st-v1_.CEL |  |  |
| iris EC | GSM524667.CEL | HG-U133_Plus_2 | E-MTAB-3732 |
| late endothelial precursor | GSM909292_S0119-135-UCB-lEPC.CEL | HG-U133_Plus_2 | E-GEOD-37045 |
|  | GSM909293_S0119-136-UCB-lEPC.CEL |  |  |
| LSEC | GSM1673671_L4_LSEC.CEL | HG-U219 | E-GEOD-68000 |
|  | GSM1673672_L8_LSEC.CEL |  |  |
|  | GSM1673673_L10_LSEC.CEL |  |  |
| lymphatic EC | GSM1528088_LEC_pop1_1.CEL | HG-U133_Plus_2 | E-GEOD-62510 |
|  | GSM1528089_LEC_pop1_2.CEL |  |  |
|  | GSM1528090_LEC_pop1_3.CEL |  |  |
|  | GSM1528091_LEC_pop2_1.CEL |  |  |
|  | GSM1528092_LEC_pop2_2.CEL |  |  |
|  | GSM1528093_LEC_pop2_3.CEL |  |  |
|  | GSM2241753_HuLEC1.CEL | HuGene-2_0-st | E-GEOD-84551 |
|  | GSM2241754_HuLEC2.CEL |  |  |
|  | GSM2241755_HuLEC3.CEL |  |  |
| placenta arterial EC | GSM1083990_female_ECA_105_5_b_3_HuGene-1_0-st-v1_.CEL | HuGene-1_0-st-v1 | E-GEOD-44368 |
|  | GSM1083991_female_ECA_127_5_a_5_HuGene-1_0-st-v1_.CEL |  |  |
|  | GSM1083992_female_ECA_135_5_a_7_HuGene-1_0-st-v1_.CEL |  |  |
|  | GSM1083993_female_ECA_84_7_b_1_HuGene-1_0-st-v1_.CEL |  |  |
|  | GSM1083999_male_ECA_117_7_b_15_HuGene-1_0-st-v1_.CEL |  |  |
|  | GSM1084000_male_ECA_92_5_b_9_HuGene-1_0-st-v1_.CEL |  |  |
|  | GSM1084001_male_ECA_98_7_b_11_HuGene-1_0-st-v1_.CEL |  |  |
|  | GSM1084002_male_ECA_100_8_b_13_HuGene-1_0-st-v1_.CEL |  |  |
| placenta venous EC | GSM1083994_female_ECV_105_6_b_4_HuGene-1_0-st-v1_.CEL | HuGene-1_0-st-v1 | E-GEOD-44368 |
|  | GSM1083995_female_ECV_127_8_b_6_HuGene-1_0-st-v1_.CEL |  |  |
|  | GSM1083996_female_ECV_135_6_b_8_HuGene-1_0-st-v1_.CEL |  |  |
|  | GSM1083997_female_ECV_84_5_b_2_HuGene-1_0-st-v1_.CEL |  |  |
|  | GSM1083998_male_ECV_100_8_b_14_HuGene-1_0-st-v1_.CEL |  |  |
|  | GSM1084003_male_ECV_117_6_c_16_HuGene-1_0-st-v1_.CEL |  |  |
|  | GSM1084004_male_ECV_92_5_c_10_HuGene-1_0-st-v1_.CEL |  |  |
|  | GSM1084005_male_ECV_98_7_b_12_HuGene-1_0-st-v1_.CEL |  |  |
| prostate EC | GSM230274.CEL | HG-U133_Plus_2 | E-GEOD-9196 |
|  | GSM230275.CEL |  |  |
|  | GSM230276.CEL |  | E-MTAB-3732 |
|  | GSM230277.CEL |  |  |
|  | GSM230278.CEL |  |  |
| pulmonary arterial EC | GSM1967875_R346_2171A.CEL | HuGene-2_0-st | E-GEOD-75793 |
|  | GSM1967876_R348_2171B.CEL |  |  |
|  | GSM1967877_R349_2171C.CEL |  |  |
|  | GSM1967878_R351_2171D.CEL |  |  |
|  | GSM1967879_R358_2171E.CEL |  |  |
|  | GSM1967880_R360_2171F.CEL |  |  |
|  | GSM1967881_R361_2171G.CEL |  |  |
|  | GSM1967882_R363_2171H.CEL |  |  |
|  | GSM1967883_R364_2171I.CEL |  |  |
|  | GSM1967884_R366_2171J.CEL |  |  |
|  | GSM1967885_R367_2171K.CEL |  |  |
|  | GSM1967886_R369_2171L.CEL |  |  |
|  | NUID-0000-0097-6604.cel | HG-U133_Plus_2 | E-MTAB-2495 |
|  | NUID-0000-0097-6607.cel |  |  |
|  | NUID-0000-0097-6609.cel |  |  |
|  | NUID-0000-0097-6629.cel |  |  |
|  | NUID-0000-0097-6632.cel |  |  |
|  | NUID-0000-0097-6635.cel |  |  |
|  | NUID-0000-0097-6638.cel |  |  |
|  | NUID-0000-0097-6652.cel |  |  |
|  | NUID-0000-0097-6655.cel |  |  |
|  | NUID-0000-0097-6657.cel |  |  |
|  | NUID-0000-0097-6662.cel |  |  |
|  | NUID-0000-0097-6675.cel |  |  |
|  | NUID-0000-0097-6679.cel |  |  |
|  | NUID-0000-0097-6681.cel |  |  |
|  | NUID-0000-0097-6685.cel |  |  |
|  | NUID-0000-0100-8017.cel |  |  |
| retina EC | GSM524664.CEL | HG-U133_Plus_2 | E-MTAB-3732 |
|  | GSM524666.CEL |  |  |
| umbilical artery EC | GSM1210811_06AN.CEL | HG-U133_Plus_2 | E-GEOD-49958 |
|  | GSM1210812_20AN.CEL |  |  |
|  | GSM1210813_39AN.CEL |  |  |
|  | GSM1210814_40AN.CEL |  |  |
|  | GSM1210815_41AN.CEL |  |  |
|  | GSM1210816_43AN.CEL |  |  |
|  | GSM1210817_06AH.CEL |  |  |
|  | GSM1210818_20AH.CEL |  |  |
|  | GSM1210819_39AH.CEL |  |  |
|  | GSM1210820_40AH.CEL |  |  |
|  | GSM1210821_41AH.CEL |  |  |
|  | GSM1210822_43AH.CEL |  |  |
| umbilical cord outgrowth EC | GSM515223.CEL | HG-U133_Plus_2 | E-MTAB-3732 |
|  | GSM515224.CEL |  |  |
|  | GSM515226.CEL |  |  |
|  | GSM515227.CEL |  |  |
|  | GSM515228.CEL |  |  |
| uterus EC | C_pool.CEL | HG-U133_Plus_2 | E-MTAB-5467 |
|  | T_pool.CEL |  |  |

**Supplementary Table 4 Transcription factor rankings**

| **Cell type** | **Module** | **TFs** | **NES** | **kME** | **RF** | **FC** |
| --- | --- | --- | --- | --- | --- | --- |
| blood outgrowth EC | Cell cycle (G1-S transition) | *E2F1 TFDP1 RB1 NFYA SP1 E2F8 NFYC* | 16.1 14 14.6 5.02 4.3 12.8 5.24 | 0.62 0.71 0.59 0.65 0.61 0.56 0.57 | 0.84 0.53 0.51 0.47 0.4 0.38 0.36 | 0.58 0.4 0.49 0.62 0.65 0.34 0.73 |
|  | Cell cycle (S-G2-M) | *E2F1 E2F8* | 14 12.7 | 0.6 0.81 | 0.69 0.38 | 0.58 0.34 |
|  | Development Focal adhesions | *DEAF1 TEAD2 ACO1* | 4.01 3.89 3.2 | 0.75 0.72 0.87 | 0.35 0.35 0.31 | 0.67 0.55 0.47 |
|  | Immune response secretion | *SPI1 IRF4* | 3.69 3.75 | 0.9 0.66 | 0.53 0.35 | 0.71 0.86 |
|  | RNA metabolism | *ELF2* | 4.35 | 0.74 | 0.43 | 1.26 |
| coronary artery EC | Cell cycle (G1-S transition) | *E2F1 SIN3A TFDP1 RB1 NFYA SP1 E2F8 NFYC* | 16.1 12.1 14 14.6 5.02 4.3 12.8 5.24 | 0.62 0.53 0.71 0.59 0.65 0.61 0.56 0.57 | 0.84 0.62 0.53 0.51 0.47 0.4 0.38 0.36 | 1.31 1.45 3.39 1.94 1.35 1.92 1.68 2.08 |
|  | Cell cycle (S-G2-M) | *E2F1 E2F8* | 14 12.7 | 0.6 0.81 | 0.69 0.38 | 1.31 1.68 |
|  | Development Focal adhesions | *DEAF1 ACO1* | 4.01 3.2 | 0.75 0.87 | 0.35 0.31 | 1.13 4.17 |
|  | Immune response (cytokine) signalling | *IRF8 BCL11A* | 3.67 4.58 | 0.84 0.53 | 0.4 0.37 | 0.55 0.68 |
|  | Immune response inflammasomes | *IRF1 IRF9 IRF5 STAT1 IRF8 STAT2* | 21.5 22.4 16.1 24.7 21.4 23.7 | 0.69 0.81 0.58 0.53 0.57 0.5 | 0.9 0.89 0.81 0.81 0.79 0.79 | 0.5 0.74 0.8 1.21 0.55 0.73 |
|  | Immune response secretion | *RELA* | 3.44 | 0.5 | 0.35 | 1.32 |
|  | RNA metabolism | *ELF2* | 4.35 | 0.74 | 0.43 | 0.73 |
| cultured HUVEC | Immune response inflammasomes | *STAT1* | 24.7 | 0.53 | 0.81 | 0.4 |
| cultured lung EC | Cell cycle (G1-S transition) | *E2F1 SIN3A TFDP1 SP1 E2F8* | 16.1 12.1 14 4.3 12.8 | 0.62 0.53 0.71 0.61 0.56 | 0.84 0.62 0.53 0.4 0.38 | 0.72 0.6 0.42 1.56 0.37 |
|  | Cell cycle (S-G2-M) | *E2F1 E2F8 BRCA1* | 14 12.7 3.24 | 0.6 0.81 0.86 | 0.69 0.38 0.34 | 0.72 0.37 0.66 |
|  | Immune response (cytokine) signalling | *IRF4 IRF8* | 6.16 3.67 | 0.53 0.84 | 0.48 0.4 | 0.9 1.32 |
|  | Immune response inflammasomes | *IRF1 IRF9 IRF5 STAT1 IRF8 STAT2 IRF7* | 21.5 22.4 16.1 24.7 21.4 23.7 15.5 | 0.69 0.81 0.58 0.53 0.57 0.5 0.83 | 0.9 0.89 0.81 0.81 0.79 0.79 0.75 | 2.35 2.03 1.21 3.72 1.32 1.96 3.5 |
| cultured umbilical artery EC | Development Focal adhesions | *DEAF1* | 4.01 | 0.75 | 0.35 | 0.56 |
|  | Immune response secretion | *SPI1 RELA* | 3.69 3.44 | 0.9 0.5 | 0.53 0.35 | 0.53 0.49 |
| dermal microvascular EC | Cell cycle (G1-S transition) | *NFYA* | 5.02 | 0.65 | 0.47 | 0.46 |
| dermal microvascular EC | Immune response (cytokine) signalling | *IRF8* | 3.67 | 0.84 | 0.4 | 3 |
|  | Immune response inflammasomes | *IRF1 IRF8* | 21.5 21.4 | 0.69 0.57 | 0.9 0.79 | 2.85 3 |
| early endothelial precursor | Immune response (cytokine) signalling | *IRF8* | 3.67 | 0.84 | 0.4 | 22.88 |
|  | Immune response inflammasomes | *IRF5 IRF8* | 16.1 21.4 | 0.58 0.57 | 0.81 0.79 | 5.33 22.88 |
| EC (unspecified) | Cell cycle (G1-S transition) | *SIN3A TFDP1 RB1 NFYA SP1 E2F8 NFYC* | 12.1 14 14.6 5.02 4.3 12.8 5.24 | 0.53 0.71 0.59 0.65 0.61 0.56 0.57 | 0.62 0.53 0.51 0.47 0.4 0.38 0.36 | 0.44 0.59 0.51 0.6 0.78 2.01 0.62 |
|  | Cell cycle (S-G2-M) | *E2F8 BRCA1* | 12.7 3.24 | 0.81 0.86 | 0.38 0.34 | 2.01 1.3 |
|  | Development Focal adhesions | *ACO1* | 3.2 | 0.87 | 0.31 | 0.47 |
|  | Immune response (cytokine) signalling | *SPI1* | 8.97 | 0.6 | 0.79 | 0.66 |
|  | Immune response inflammasomes | *IRF1 IRF9 IRF5 IRF7* | 21.5 22.4 16.1 15.5 | 0.69 0.81 0.58 0.83 | 0.9 0.89 0.81 0.75 | 1.23 1.89 1.14 2.22 |
|  | Immune response secretion | *SPI1 RELA* | 3.69 3.44 | 0.9 0.5 | 0.53 0.35 | 0.66 0.66 |
|  | RNA metabolism | *ELF2* | 4.35 | 0.74 | 0.43 | 1.39 |
| endothelial precursor | Cell cycle (G1-S transition) | *E2F1 E2F8* | 16.1 12.8 | 0.62 0.56 | 0.84 0.38 | 0.63 0.55 |
|  | Cell cycle (S-G2-M) | *E2F1 E2F8* | 14 12.7 | 0.6 0.81 | 0.69 0.38 | 0.63 0.55 |
|  | Immune response (cytokine) signalling | *SPI1 IRF4 IRF8 BCL11A* | 8.97 6.16 3.67 4.58 | 0.6 0.53 0.84 0.53 | 0.79 0.48 0.4 0.37 | 5.22 1.32 5.98 1.83 |
|  | Immune response inflammasomes | *IRF1 IRF9 IRF5 STAT1 IRF8 IRF7* | 21.5 22.4 16.1 24.7 21.4 15.5 | 0.69 0.81 0.58 0.53 0.57 0.83 | 0.9 0.89 0.81 0.81 0.79 0.75 | 2.32 2.02 1.86 2.58 5.98 2.75 |
|  | Immune response secretion | *SPI1 IRF4* | 3.69 3.75 | 0.9 0.66 | 0.53 0.35 | 5.22 1.32 |
| fresh HUVEC | Cell cycle (G1-S transition) | *SP1 NFYC* | 4.3 5.24 | 0.61 0.57 | 0.4 0.36 | 0.5 0.44 |
|  | RNA metabolism | *ELF2* | 4.35 | 0.74 | 0.43 | 1.89 |
| fresh umbilical artery EC | Cell cycle (G1-S transition) | *NFYC* | 5.24 | 0.57 | 0.36 | 0.51 |
|  | Development Focal adhesions | *ACO1* | 3.2 | 0.87 | 0.31 | 0.32 |
|  | Immune response inflammasomes | *IRF9 IRF7* | 22.4 15.5 | 0.81 0.83 | 0.89 0.75 | 2.76 2.19 |
|  | Immune response secretion | *RELA* | 3.44 | 0.5 | 0.35 | 0.49 |
|  | RNA metabolism | *ELF2* | 4.35 | 0.74 | 0.43 | 2.3 |
| immortalised microvascular EC | Cell cycle (G1-S transition) | *E2F8* | 12.8 | 0.56 | 0.38 | 2.75 |
|  | Cell cycle (S-G2-M) | *E2F8 BRCA1* | 12.7 3.24 | 0.81 0.86 | 0.38 0.34 | 2.75 2.41 |
|  | Immune response (cytokine) signalling | *IRF8* | 3.67 | 0.84 | 0.4 | 2.77 |
|  | Immune response inflammasomes | *STAT1 IRF8* | 24.7 21.4 | 0.53 0.57 | 0.81 0.79 | 0.52 2.77 |
|  | RNA metabolism | *ELF2* | 4.35 | 0.74 | 0.43 | 1.52 |
| intestinal lymphatic EC | Cell cycle (S-G2-M) | *BRCA1* | 3.24 | 0.86 | 0.34 | 0.52 |
|  | Development Focal adhesions | *DEAF1 TEAD2* | 4.01 3.89 | 0.75 0.72 | 0.35 0.35 | 1.53 3.8 |
|  | Immune response (cytokine) signalling | *SPI1* | 8.97 | 0.6 | 0.79 | 1.48 |
| intestinal lymphatic EC | Immune response secretion | *SPI1 RELA* | 3.69 3.44 | 0.9 0.5 | 0.53 0.35 | 1.48 1.51 |
| LSEC | Cell cycle (S-G2-M) | *E2F8* | 12.7 | 0.81 | 0.38 | 0.2 |
|  | Development Focal adhesions | *TEAD2* | 3.89 | 0.72 | 0.35 | 4.42 |
|  | Immune response (cytokine) signalling | *SPI1 IRF4 IRF8 BCL11A* | 8.97 6.16 3.67 4.58 | 0.6 0.53 0.84 0.53 | 0.79 0.48 0.4 0.37 | 5.6 1.86 2.25 0.54 |
|  | Immune response inflammasomes | *IRF9 IRF8 STAT2 IRF7* | 22.4 21.4 23.7 15.5 | 0.81 0.57 0.5 0.83 | 0.89 0.79 0.79 0.75 | 2.55 2.25 4.95 2.74 |
|  | Immune response secretion | *SPI1 IRF4 RELA* | 3.69 3.75 3.44 | 0.9 0.66 0.5 | 0.53 0.35 0.35 | 5.6 1.86 2.85 |
| placenta arterial EC | Cell cycle (G1-S transition) | *E2F1 TFDP1 RB1 NFYA E2F8* | 16.1 14 14.6 5.02 12.8 | 0.62 0.71 0.59 0.65 0.56 | 0.84 0.53 0.51 0.47 0.38 | 1.32 2.44 2.1 2.19 2.49 |
|  | Cell cycle (S-G2-M) | *E2F1 E2F8* | 14 12.7 | 0.6 0.81 | 0.69 0.38 | 1.32 2.49 |
|  | Development Focal adhesions | *DEAF1 TEAD2 ACO1* | 4.01 3.89 3.2 | 0.75 0.72 0.87 | 0.35 0.35 0.31 | 1.48 2.17 2.42 |
|  | Immune response (cytokine) signalling | *SPI1 IRF8* | 8.97 3.67 | 0.6 0.84 | 0.79 0.4 | 1.49 0.57 |
|  | Immune response inflammasomes | *IRF1 IRF8* | 21.5 21.4 | 0.69 0.57 | 0.9 0.79 | 0.56 0.57 |
|  | Immune response secretion | *SPI1 RELA* | 3.69 3.44 | 0.9 0.5 | 0.53 0.35 | 1.49 1.89 |
|  | RNA metabolism | *ELF2* | 4.35 | 0.74 | 0.43 | 0.68 |
| placenta venous EC | Cell cycle (G1-S transition) | *TFDP1 NFYC* | 14 5.24 | 0.71 0.57 | 0.53 0.36 | 2.51 2.03 |
|  | Development Focal adhesions | *DEAF1 ACO1* | 4.01 3.2 | 0.75 0.87 | 0.35 0.31 | 1.4 3.09 |
|  | Immune response inflammasomes | *IRF8 STAT2* | 21.4 23.7 | 0.57 0.5 | 0.79 0.79 | 0.56 1.46 |
|  | Immune response secretion | *SPI1* | 3.69 | 0.9 | 0.53 | 1.66 |
|  | RNA metabolism | *ELF2* | 4.35 | 0.74 | 0.43 | 0.7 |
| prostate EC | Cell cycle (G1-S transition) | *TFDP1 SP1 E2F8 NFYC* | 14 4.3 12.8 5.24 | 0.71 0.61 0.56 0.57 | 0.53 0.4 0.38 0.36 | 0.33 0.5 0.22 0.45 |
|  | Cell cycle (S-G2-M) | *E2F8 BRCA1* | 12.7 3.24 | 0.81 0.86 | 0.38 0.34 | 0.22 0.56 |
|  | Development Focal adhesions | *DEAF1 TEAD2 ACO1* | 4.01 3.89 3.2 | 0.75 0.72 0.87 | 0.35 0.35 0.31 | 0.62 0.48 0.35 |
|  | Immune response (cytokine) signalling | *IRF4 IRF8 BCL11A* | 6.16 3.67 4.58 | 0.53 0.84 0.53 | 0.48 0.4 0.37 | 1.31 10.45 1.43 |
|  | Immune response inflammasomes | *IRF1 IRF5 STAT1 IRF8 IRF7* | 21.5 16.1 24.7 21.4 15.5 | 0.69 0.58 0.53 0.57 0.83 | 0.9 0.81 0.81 0.79 0.75 | 9.29 1.55 0.44 10.45 2.71 |
|  | Immune response secretion | *IRF4* | 3.75 | 0.66 | 0.35 | 1.31 |
|  | RNA metabolism | *ELF2* | 4.35 | 0.74 | 0.43 | 1.77 |
| pulmonary arterial EC | Cell cycle (S-G2-M) | *E2F1 E2F8* | 14 12.7 | 0.6 0.81 | 0.69 0.38 | 0.77 0.67 |
| umbilical artery EC | Cell cycle (G1-S transition) | *SIN3A E2F8* | 12.1 12.8 | 0.53 0.56 | 0.62 0.38 | 0.49 2.92 |
|  | Cell cycle (S-G2-M) | *E2F8 BRCA1* | 12.7 3.24 | 0.81 0.86 | 0.38 0.34 | 2.92 2.14 |
|  | Development Focal adhesions | *ACO1* | 3.2 | 0.87 | 0.31 | 0.39 |
|  | Immune response (cytokine) signalling | *IRF4* | 6.16 | 0.53 | 0.48 | 0.91 |
|  | Immune response inflammasomes | *STAT1 IRF7* | 24.7 15.5 | 0.53 0.83 | 0.81 0.75 | 0.42 0.65 |
|  | Immune response secretion | *IRF4 RELA* | 3.75 3.44 | 0.66 0.5 | 0.35 0.35 | 0.91 0.58 |
|  | RNA metabolism | *ELF2* | 4.35 | 0.74 | 0.43 | 1.38 |
| (NES: maximal network enrichment score ; kME: module eigengene ; RF: Regulated fraction of the respective module ; FC: fold change) | | | | | | |
